# Identifying novel radioprotective drugs via salivary gland tissue chip screening

**DOI:** 10.1101/2023.07.12.548707

**Authors:** Lindsay Piraino, Chiao Yun Chen, Jared Mereness, P. M. Dunman, Catherine Ovitt, Danielle Benoit, Lisa DeLouise

## Abstract

During head and neck cancer treatment, off-target ionizing radiation damage to the salivary glands commonly causes a permanent loss of secretory function. Due to the resulting decrease in saliva production, patients have trouble eating, speaking and are predisposed to oral infections and tooth decay. While the radioprotective antioxidant drug Amifostine is approved to prevent radiation-induced hyposalivation, it has intolerable side effects that limit its use, motivating the discovery of alternative therapeutics. To address this issue, we previously developed a salivary gland mimetic (SGm) tissue chip platform. Here, we leverage this SGm tissue chip for high-content drug discovery. First, we developed in-chip assays to quantify glutathione and cellular senescence (β-galactosidase), which are biomarkers of radiation damage, and we validated radioprotection using WR-1065, the active form of Amifostine. Following validation, we tested other reported radioprotective drugs, including, Edaravone, Tempol, N-acetylcysteine (NAC), Rapamycin, Ex-Rad, and Palifermin, confirming that all drugs but NAC and Ex-Rad exhibited robust radioprotection. Next, a Selleck Chemicals library of 438 FDA-approved drugs was screened for radioprotection. We discovered 25 hits, with most of the drugs identified with mechanisms of action other than antioxidant activity. Hits were down-selected using EC_50_ values and pharmacokinetics and pharmacodynamics data from the PubChem database leading to testing of Phenylbutazone (anti-inflammatory), Enoxacin (antibiotic), and Doripenem (antibiotic) for *in vivo* radioprotection in mice using retroductal injections. Results confirm that Phenylbutazone and Enoxacin exhibited equivalent radioprotection to Amifostine. This body of work demonstrates the development and validation of assays using a SGm tissue chip platform for high-content drug screening and the successful *in vitro* discovery and *in vivo* validation of novel radioprotective drugs with nonantioxidant primary indications pointing to possible, yet unknown novel mechanisms of radioprotection.

## 1. Introduction

During head and neck cancer treatment, ionizing radiation damage to the salivary glands often causes a permanent loss of secretory function and reduced salivary flow. Due to decreased saliva production, patients have trouble eating, speaking, and swallowing^1, 2^. Additionally, patients are at an increased risk of oral infections and tooth decay and suffer a reduced quality of life^2, 3^. Current treatment options, including sialogogues, mouthwashes, and chewing gum, only provide temporary relief, and there is no cure^2^. Several strategies have been proposed to alleviate this damage, including cell transplantation^4–7^ and gene therapy^8–11^. Despite promising results, these methods have remained experimental and are targeted toward patients already experiencing xerostomia. Hence, there is an unmet need to provide current and future head and neck cancer patients with preventative therapies to protect salivary gland function.

Intensity-modulated radiation therapy (IMRT) and the radioprotective drug Amifostine are used clinically to prevent salivary gland damage. IMRT involves using 3D imaging to target the radiation beams at the tumor and away from sensitive organs such as the salivary gland^12^. While this method can be beneficial in some cases, there are mixed results on patient-reported claims of dry mouth. Furthermore, IMRT is sometimes impossible due to tumor location^12, 13^. The antioxidant Amifostine is the only FDA-approved drug to prevent radiation-induced xerostomia. However, its use is often discontinued during fractionated radiation regimens due to severe side effects, including nausea, vomiting, and hypotension^14, 15^. Additionally, its short half-life *in vivo* limits its efficacy, as the drug is cleared within minutes of administration^16^. These drawbacks highlight the critical need to discover new radioprotective drugs to prevent xerostomia.

To address this need, we developed a microbubble (MB) array-based tissue chip consisting of 3D salivary gland tissue mimetics (SGm) entrapped within a matrix metalloproteinase degradable poly(ethylene glycol) (PEG)-based hydrogel engineered extracellular matrix (eECM) that maintains secretory behavior^17^. The spherical architecture of the MB combined with the cellularly degradable eECM creates a distinct niche that promotes cell viability and maintenance of the secretory acinar cells based on gene and protein expression as well as secretory function based on tissue architecture, gene expression, and secretory agonist-responsive calcium signaling^17^. In addition, this platform was validated for use in radioprotection studies using immunohistochemical staining of individual SGm to quantitate foci of DNA damage markers γH2AX and 53BP1 after irradiation. Analysis of control chips versus chips treated with WR1065, the active form of Amifostine, showed a reduction in DNA damage with drug treatment^17^.

Array-based assays that can interrogate each SGm on the chip (∼280 MBs/cm^2^) are necessary to enable high-content drug screening. Based on literature, several assays commonly used to report radiation-induced cellular damage were tested in the tissue chip format. Assays tested included reactive oxygen species (ROS), apoptosis, secretion, and cytotoxicity at various time points post-radiation (**Table S.1**). Based on signal-to-noise ratio and reproducibility, the glutathione ^18^ and cellular senescence^19^ assays were selected for further development for high-content screening of radioprotective drugs.

Assays were tested with 0 Gy, 15 Gy, and 15 Gy + 4 mM WR1065 to confirm usefulness for measuring radiation damage and radioprotection of individual SGm within the MB-hydrogel tissue chip (40-50 MBs per chip). The assays were further tested with other reported radioprotective drugs, including Tempol^20, 21^, N-acetylcysteine^22^, Edaravone^23^, Rapamycin^24^, Ex-Rad^25, 26^, and Palifermin^27, 28^. Next a library of 438 FDA-approved drugs was screened for radioprotection using both assays, identifying 25 double hits that were further investigated for suitability as potential therapeutics informed by data within the PubChem database and experimentally determined dose-response relationships. Three of the hits, Phenylbutazone (anti-inflammatory), Enoxacin (antibiotic), and Doripenem (antibiotic), were chosen for *in vivo* radioprotection testing of the mouse submandibular gland using retroductal injections. Results confirm that Phenylbutazone and Enoxacin exhibited equivalent radioprotection to Amifostine.

## 2. Materials and Methods

### Materials

Detailed information on the drugs used for assay development and validation, including, Tempol, N-acetylcysteine, Edaravone, WR1065, Rapamycin, Ex-Rad, and Palifermin is listed in **Table S.2**. Drug screening was completed using a 438 Selleck Chemicals library of FDA-approved drugs (**Table S.3)**. The screen’s 25 top drug hits **(Table S.4)** were purchased from Selleck Chemicals for dose-response studies and prepared and stored per manufacturer’s instructions.

### Animals

Female SKH1 hairless mice, backcrossed 6 generations with C57BL/6J mice, aged 6-12 weeks were used in this study for in vitro assay development and drug discovery. Female C57BL/6J mice age 6-8 weeks were used for in vivo validation studies. Only female mice were used due to known sex differences in rodent salivary glands, with female glands more accurately emulating human salivary gland structure and function^29, 30^. Animals were maintained on a 12 hr light/dark cycle and group-housed with food and water available *ad libitum*. All procedures were approved and conducted per the University Committee on Animal Resources at the University of Rochester Medical Center (UCAR #2010-024E, UCAR-2008-016E).

### Microbubble (MB) array fabrication

Microbubble (MB) arrays were fabricated in poly(dimethyl) siloxane (PDMS) using gas expansion molding as previously described^17, 31, 32^. PDMS (Dow Corning Sylgard 184) was mixed in a 10:1 base-to-curing agent ratio and poured over a silicon wafer template consisting of deep etched cylindrical pits with a 200 µm diameter, spaced 600 µm apart on a square lattice. The PDMS was cured at 100 °C for 2 hrs before peeling off the template, resulting in an array of spherical cavities with 200 µm opening and ∼350 µm diameter. Circular chips with 0.7 cm diameter (48 well plates) or 0.5 cm diameter (96 well plates) were punched from the PDMS cast and glued into well plates using a 5:1 ratio of PDMS cured at 60 °C for 8 hours. The MB arrays were primed in a desktop vacuum chamber with 70% ethanol to facilitate air removal from the MBs and replacement with fluid. Ethanol was exchanged for PBS and arrays were incubated overnight before cell seeding.

### Cell isolation

Mice were euthanized and the submandibular glands were removed and chopped with a razor blade for 5 min. The tissue was then incubated in Hank’s buffered salt solution (HBSS) containing 15 mM HEPES, 50 U/mL collagenase type II (Thermo Fisher 17101015), and 100 U/mL hyaluronidase (Sigma Aldrich H3506) at 37 °C for 30 min. Cells were centrifuged, resuspended in HBSS with 15 mM HEPES, and passed through 100 µm and 20 μm mesh filters to isolate clusters between 20-100 µm. The digestion protocol produces cell cluster sizes evenly distributed between 20 to 100 µm^33^. As described below, the isolated clusters were combined with hydrogel precursor solution and seeded in MB array chips.

### MB-hydrogel encapsulation of salivary gland cells

Isolated submandibular gland cell clusters (20-100 µm) were encapsulated within matrix-metalloproteinase (MMP) degradable poly(ethylene glycol) (PEG) hydrogels within MB arrays (**Fig. 1**) as previously described^17^. Briefly, the cells (**Fig. 1A**) were resuspended in hydrogel precursor (**Fig. 1B**) solution containing 2 mM norbornene-functionalized 4-arm 20 kDa PEG-amine macromers, 4 mM of the dicysteine functionalized MMP degradable peptide (GKKCGPQG↓IWGQCKKG), 0.05 wt% of the photoinitiator lithium phenyl-2,4,6-trimethylbenzoylphosphinate (LAP)^34^, and 0.1 mg/mL laminin in PBS^4, 5, 17, 35^. The cell/gel precursor solution (25 µL for 48 wells; 20 µL for 96 wells) was pipetted onto the MB chips and incubated for 30 min, pipetting every 10 min to redisperse cells that had settled onto the surface of the chip. The hydrogels were polymerized *in situ* using a Hand-Foot 1000 A broad spectrum UV light (UVA: 5 mW/cm^2^; UVB: 0.4 mW/cm^2^) with a UVC filter for 1.5 min and cultured with media (0.5 mL for 48 well; 150 µL for 96 well), with media changes every 2 days. Culture medium consisted of Dulbecco’s Modified Eagle Medium (DMEM):Ham’s F-12 Nutrient Mixture (1:1) supplemented with 100 U/mL Penicillin and 100 μg/mL Streptomycin, 2 mM Glutamine, 0.5x N2 supplement, 2.6 ng/mL insulin, 2 nM dexamethasone, 20 ng/mL epidermal growth factor (EGF), and 20 ng/mL basic fibroblast growth factor (bFGF).

**Figure 1:**
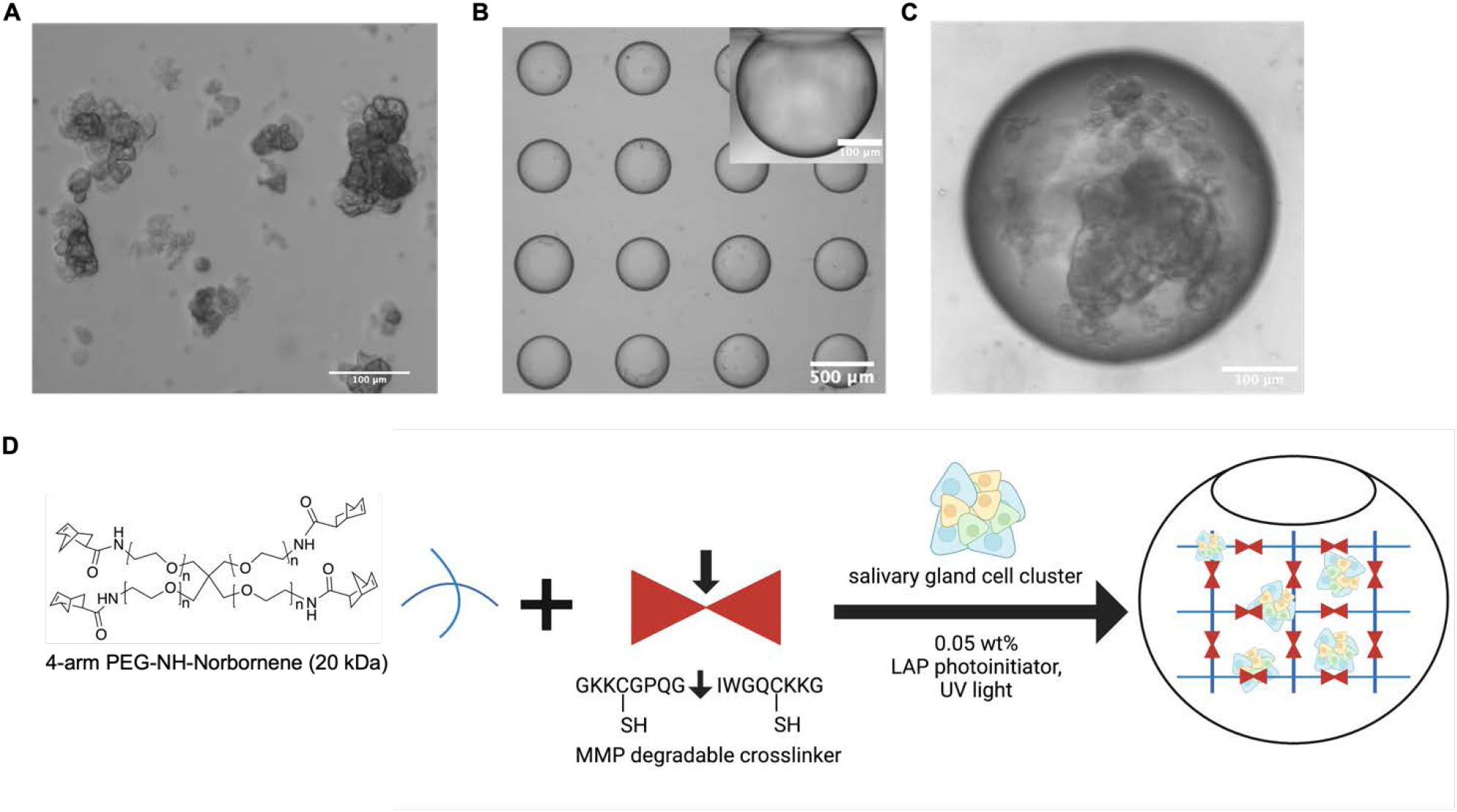
Overview of the salivary gland tissue chip. Isolated primary salivary gland cell clusters 20-100 µm (A). Clusters are seeded into the MB arrays. Inset image showing the cross-sectional view of a MB (B). Clusters over time aggregate and grow to form spheres within the MBs, example shown at day 7 (C). Schematic representation of hydrogel encapsulation of salivary gland cell clusters within an MB chip created using Biorender.com (license #AD25LJLLID) (D).

### Glutathione assay

A glutathione assay was developed for in-chip measurements by adapting the Cellular Glutathione Detection Assay Kit (Cell Signaling Technology #13859). The monochlorobimane reagent was prepared by reconstitution in DMSO per manufacturer’s directions. For 96 well plates, 10 µL of prepared reagent (1:50 ratio of monochlorobimane (MCB) to Tris assay buffer, per manufacturer’s instructions) was added to wells containing 100 µL of culture media and incubated for 30 min at 37 °C, washed with PBS, and imaged using an Olympus IX70 microscope with a DAPI filter (Excitation/Emission: 358 nm/461 nm).

### Senescence assay

A cellular senescence assay for the chips was developed by adapting the Cellular Senescence Detection Kit – SPiDER-βGal (Dojindo Molecular Technologies, Inc SG04). Balifomycin A1 and SPiDER-βGal stock solutions were prepared in DMSO per manufacturer’s directions. The assay was performed by first incubating the chips with Balifomycin A1 (1:1000 dilution in media) for 1 hr at 37 °C. The solution was removed, replaced with 30 µL of media containing Balifomycin A1 (1:1000 dilution) and SPiDER-βGal (1:500 dilution), and incubated for 45 min at 37 °C. Chips were washed twice with media and imaged using a fluorescence microscope with a Texas Red filter (Excitation/Emission: 580 nm/604 nm).

### Drug mechanism meta-analysis

Drug interaction data was examined using PubChem BioAssays results for the hits discovered in the Selleck Chemicals drug library screen described below ^36–38^. Pathway analysis was performed using the Drug Set Enrichment Analysis (DSEA) tool to identify pathways with gene expression patterns significantly impacted by treatment with the drug hits (p<0.05) ^39^.

### Image quantification and statistical analysis

For the glutathione and senescence assays, images were quantified in ImageJ. Regions of interest (ROIs) were created by thresholding on the fluorescence signal (localized to the SGm), and each ROI’s mean intensity was measured. Data were graphed and statistical analyses (ANOVA with Tukey’s post-hoc test) were performed using GraphPad Prism 9. Schematic diagrams were created using Biorender (biorender.com) under institutional site license, agreement number VP25LG6DJ3.

### Drug treatment and irradiation

For radioprotection experiments, SGm were cultured in MB-hydrogel chips for 4 days, then drugs were added to the chips 30 min before radiation and washed out with media 30 min post-radiation to parallel how Amifostine is used clinically^40^. A dose of 15 Gy ionizing radiation was delivered using a JL Shepherd ^137^Cs irradiator. Drug treatment schemes and radiation doses were established in our previous work^17^. The glutathione and senescence assays were performed at 4-and 5-days post-radiation. For assay validation experiments, at least 3 chips (N = 3) were used for each drug, corresponding to > 100 MBs (n > 100); these values are listed in the figure captions for each experiment. Mean, standard deviation, and statistics were calculated based on the number of MBs (n).

The same treatment scheme was used for screening the Selleck Chemicals library, with drugs administered at 100 µM. One MB chip (N = 1) was used per drug, with ∼40-50 MBs per chip (n = 40-50); statistics were calculated using the number of MBs (n) and compared to 0 Gy controls. Drugs were first screened using the glutathione assay. Drugs that exhibited statistically insignificant differences compared to the 0 Gy control (hits) with the glutathione assay were then tested with the senescence assay to discover double hits. The glutathione assay was selected as the first screen because it had higher signal-to-noise ratio, more rapid throughput (30 mins versus 2 hrs for completion of the senescent assay), lower assay kit cost, and a higher shelf-life after kit reconstitution.

### Retroductal injection and irradiation

Retroductal injection delivery of drugs to the murine submandibular gland has been described in detail^41, 42^. Briefly, 6-10 week old female C57Bl/6J mice were anesthetized by intraperitoneal injection of sterile saline solution of 100 mg/kg ketamine and 10 mg/kg xylazine. Maxillary incisors were secured over a metal beam, while an elastic band provided tension from behind the mandibular incisors. The mouth was widened using a custom steel retractor to apply pressure to the buccal mucosa and the tongue was retracted and cotton placed in the oral cavity. The wire inset of a 32G intracranial catheter was cut at 45° to create a bevel. The beveled wire created a shallow puncture in the left salivary papilla. A beveled catheter section containing the wire insert for support was gently inserted into the puncture site. The catheter was removed, and 1 mg/kg atropine was administered by intraperitoneal injection. After 10 min, the needle of a Hamilton syringe, loaded with vehicle or drug solution, was inserted into the catheter, and the catheter was inserted into the orifice produced in the papilla. Drug solutions were injected by hand at 10 µl/min at a volume of 1 µl/g of body weight. Following injection, the pressure was maintained on the syringe for 1 min to ensure material retention before removal of the catheter. The cotton ^43^and retractor were removed from the oral cavity, and the elastic band and metal beam were released from the incisors. The known radioprotectant, WR-1065 (50 mg/kg, saline) was used as a control for radioprotection. Phenylbutazone, Enoxacin, and Doripenem hydrate were treated at 50, 0.3, and 26 mg/kg (N = 4-6 per group). Drug doses were chosen based on solubility and published doses for in vivo studies^44, 45^. Saline was used as the vehicle for all compounds, except Phenylbutazone, which required corn oil due to poor aqueous solubility.

Mice injected with vehicle controls (saline or corn oil) or drug were treated with 0 Gy or 15 Gy within 15-30 min of injection, the submandibular glands were irradiated as described previously ^46–48^. The head and neck region was positioned over the slit of a custom collimator, which allowed body shielding. Mouse submandibular glands were exposed to a single dose of 15 Gy gamma radiation delivered by a ^137^Cs radiation source. Prior studies demonstrate that this single dose causes xerostomia in mice equivalent to fractionated dosing that recapitulates human sequalae of salivary gland radiation damage ^17, 49^. Animals were allowed to recover for 48 hours to measure persistent long-lived DNA damage, after which, the submandibular glands were harvested, fixed, sectioned, and analyzed using immunohistochemistry.

### Immunohistochemical analysis

Submandibular glands and SGm were isolated and fixed in 4% paraformaldehyde overnight at 4 °C. Tissues were paraffin-embedded and then cut into 5 μm sections. Slides were treated with HIER buffer (10 mM sodium citrate, 0.05% Tween-20, pH 6.0) for antigen retrieval in a pressure cooker for 10 minutes then sections were blocked in CAS-block histochemical reagent (Thermo Fisher Scientific, 008120). Permeabilization was performed with 0.5% Triton X-100 in PBS for 5 minutes. Immunostaining was performed overnight (at 4 °C) with primary antibody for γH2AX (EMD Millipore, 05-636). Alexa-Fluor 594-conjugated donkey anti-mouse IgG was diluted 1:500 (Invitrogen, A21203) as secondary antibody and applied on sections for 1 hour at room temperature. Following a PBS rinse, 10 μg/ml DAPI (Invitrogen, Carlsbad, CA) in PBS was applied to sections for 5 minutes. Sections were washed thrice in PBS for 5 minutes and the slides were mounted using Immu-Mount mounting medium (Thermo). Microscopic images were acquired using a Leica TCS SP5 confocal microscope with a 100X oil immersion objective and Argon laser. Analysis of images was performed in ImageJ.

## 3. Results and Discussion

### Salivary Gland Tissue Chip

Our previously reported salivary gland tissue chip^17^ was leveraged for high-content drug screening to identify novel radioprotective compounds. The chip platform consists of an array of near-spherical microbubble (MB) cavities formed in poly(dimethylsiloxane) (PDMS). Each chip, containing ∼50 MBs, affixed within wells of a 96-well plate (**Fig. 1C**). Primary salivary gland cell clusters (**Fig. 1A**), suspended in a poly(ethylene glycol) (PEG) hydrogel precursor and MMP-degradable peptide crosslinker solution together with the photoinitiator LAP, were applied to the chip allowing the clusters to deposit into the MBs **(Fig. 1D)**. *In situ* polymerization of the hydrogel was achieved using long-wave, low intensity UV light. Over time, the cell clusters aggregate and proliferate to form SGm **(Fig. 1C**).

### High-throughput methods to assess drug radioprotection

In prior studies^17^, immunohistochemical (IHC) staining was used to quantify the number γH2AX and 53BP1 puncta within nuclei, which are sensitive markers for double-strand breaks^50^ and, thus, direct measures of radiation damage/protection. IHC staining is, however, a laborious time and resource-consuming process requiring retrieval of the SGm from the MB array chip, tissue sectioning, staining, and imaging. While IHC enables sensitive analyses of radiation damage, it completely abrogates the goal of in situ high-content screening for which the tissue chip was developed. As mentioned above, several assays commonly used to assess radiation-induced cellular damage were tested in the tissue chip format at various time points post-radiation (**Table S.1**). Based on signal-to-noise ratio and reproducibility, the glutathione ^18^ and senescence^19^ assays were selected for further development. The glutathione assay was used to measure glutathione levels at various time points post-radiation to determine the optimal time for detecting differences between 0 Gy and 15 Gy. Based on the data, the greatest signal separation was measured at 4 days post-irradiation (**Fig. S1)**. This time point is similar to previous reports on decreases in glutathione post-radiation^18^ and was used for all experiments moving forward.

Next, WR1065 was tested to identify effective concentration(s) that prevented changes in glutathione levels post-radiation. A SGm tissue chip was cultured for 4 days to allow spheres to form^17^. The chip was then treated with WR1065 30 min before and during radiation, followed by drug wash out with media 30 min post-radiation (**Fig. 2A**). This dosing scheme is consistent with the use of Amifostine clinically^14^ and with our previous work^17^. The glutathione assay was performed 4 days post-radiation (**Fig. 2A**). Example images show high levels of glutathione at 0 Gy (**Fig. 2B**) that is decreased by 15 Gy radiation (**Fig. 2C**) and maintained with 4 mM WR1065 (**Fig. 2D**). Quantification shows that 0.1 mM and 0.4 mM WR1065 were ineffective at preventing radiation damage. In contrast, 1 mM and 4 mM WR1065 provided significant protection (**Fig. 2E**). The 4 mM dose corresponds with our previous work on DNA damage markers γH2AX and 53BP1^17^ and values from literature^51^ and establishes the range of effective concentrations for WR1065 treatment *in vitro.* Moreover, it is clinically relevant as 15-30 minutes before radiation, patients are administered Amifostine intravenously at 200 mg/m^2^ ^15, 40^. Assuming an average adult body surface area of 17,000 cm^2^ and a blood volume of 1.35 gallons (5.1 liters)^52, 53^, Amifostine is administered at 300 µM.

**Figure 2:**
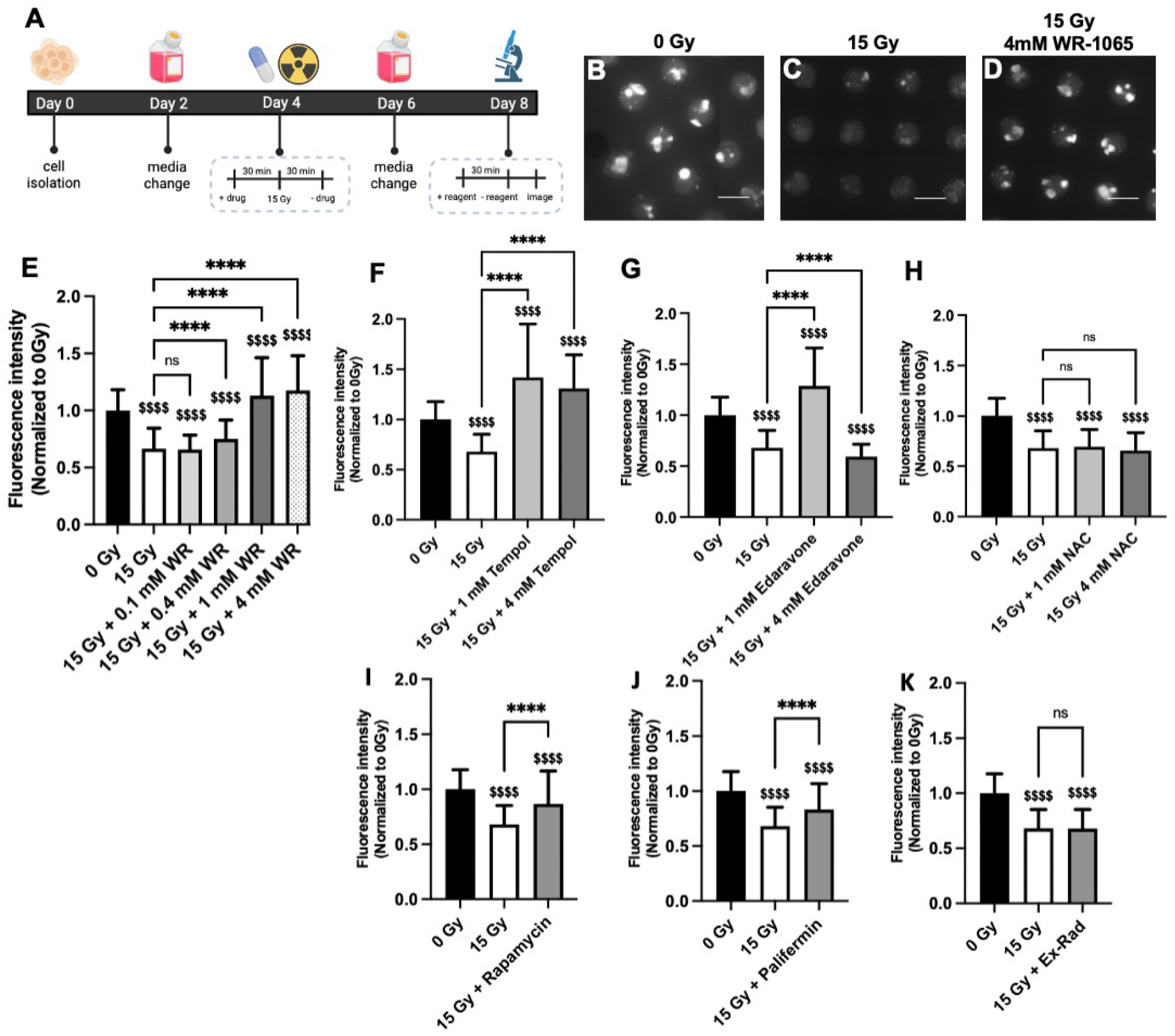
Glutathione levels indicated a dose-response to radioprotective drug, WR1065 and other reported radioprotective compounds. Timeline of drug treatment and assay for glutathione at 4 days post-radiation created using Biorender.com (A). Representative images of the glutathione assay for 0 Gy (B), 15 Gy (C), 15 Gy + 4 mM WR1065 (WR) (D) and quantification of the fluorescence intensity of individual MBs normalized to 0 Gy (E). Brackets with asterisks compared to 15 Gy: ns = nonsignificant, **** = p<0.0001; Money sign compared to 0 Gy: $$$$ = p<0.0001; WR = WR1065; N (# of chips) ≥ 3, n (# of MBs) ≥ 150. Scale bar is 600 µm. Drugs are used to treat SGm on day 4 for 30 min prior to and 30 min after radiation and senescence is analyzed on day 9 for Tempol (F), Edaravone (G), N-acetylcysteine (H), 50 µM Rapamycin (I), 100 ng/mL Palifermin (J), and 50 µM Ex-Rad (K). Brackets with asterisks compared to 15 Gy: **** = p<0.0001; Money sign compared to 0 Gy: $$$$ = p<0.0001, $$$ = p<0.001, $$ = p<0.01, ns = nonsignificant; NAC = N-acetylcysteine; N (# of chips) ≥ 3, n (# of MBs) ≥ 120.

Since WR1065 is an antioxidant and mediates radioprotective effects through free radical scavenging and induction of superoxide dismutase expression^54^, we tested other antioxidants implicated as radioprotective (Tempol, Edaravone, N-acetylcysteine). Tempol exhibited excellent radioprotection at both 1 and 4 mM (**Fig. 2F**) which is consistent with prior studies ^20, 49, 55^. Edaravone showed complete radioprotection at 1 mM but not at 4 mM (**Fig. 2G**). Edaravone maintained ∼43% of glutathione levels at 0.1 mM (**Fig. S2A)**, suggesting that the optimal concentration range for Edaravone might be lower than WR1065, which is suggested by literature indicating radio-protective dose ranges are 100-1000 µM^23, 56, 57^. For N-acetylcysteine (NAC), no improvement in glutathione levels was observed at 1 or 4 mM compared to untreated SGm **(Fig. 2H);** however, glutathione levels were rescued by 56% when treated with 10 mM NAC (**Fig. S2B)**, consistent with literature^22^.

Drugs reported with non-antioxidant radioprotective mechanisms were also tested using the glutathione assay. Rapamycin is an mTOR inhibitor reported to restore salivary flow rate post-irradiation in swine^24^. Ex-Rad reduces p53-dependent and independent apoptosis^25^. Palifermin (keratinocyte growth factor) has been reported to stimulate salivary gland stem/progenitor cell expansion post-radiation^27^. Using drug concentrations based on literature^26, 28^, our data shows some protection against glutathione level changes resulting from radiation with 50 µM rapamycin (58%, **Fig. 2I**) and 100 ng/mL Palifermin (47%, **Fig. 2J)**, but no protection from 50 µM Ex-Rad (**Fig. 2K**).

For senescence, a protocol similar to the glutathione assay was followed to determine the optimal time point post-radiation for detecting a change in senescence between 0 Gy, 15 Gy, as measured by senescence-associated β-galactosidase activity. A time point of 5 days post-radiation was optimal (**Fig. S3**), similar to a previous study ^19^. WR1065 radioprotection was tested by adding drug to the chips 30 min before radiation followed by wash out 30 min post-radiation (**Fig. 3A**). An expected increase in senescence was detected for SGm exposed to 15 Gy compared to 0 Gy (**Figs. 3B,C**), which was restored to levels equivalent to 0 Gy with the addition of WR1065 (**Fig. 3D**). Quantification shows that both 1 mM and 4 mM WR1065 treatment resulted in complete radioprotection (**Fig. 3E**), consistent with the glutathione assay results.

**Figure 3:**
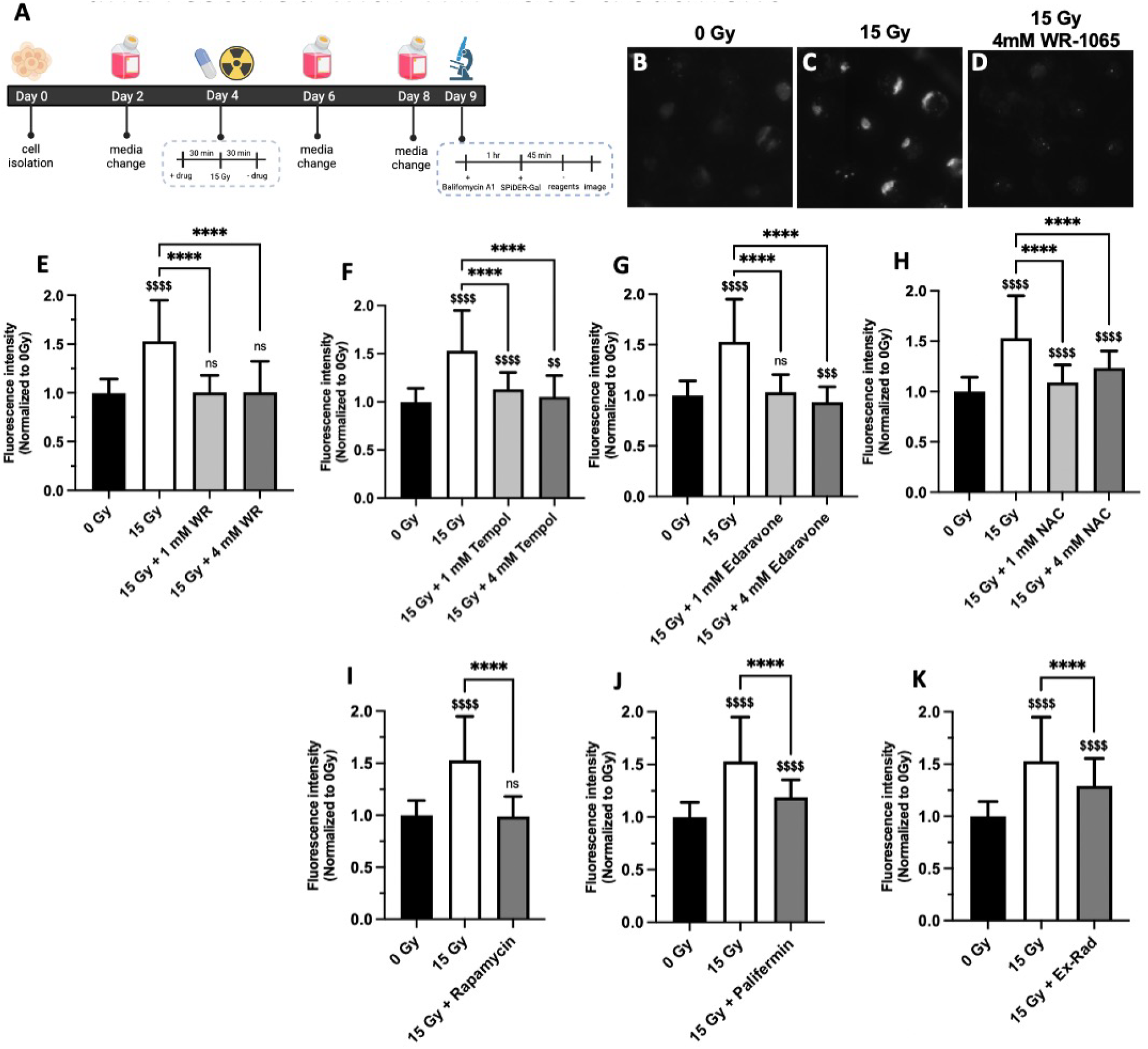
Senescence is increased with radiation and restored to unirradiated control levels with WR-1065 and other reported radioprotective compounds. Timeline of drug treatment and assay for senescence at 5 days post-radiation created using Biorender.com (A). Representative images of the senescence assay for 0 Gy (B), 15 Gy (C), 15 Gy + 4 mM WR-1065 (D) and quantification of the fluorescence intensity of individual MBs normalized to 0 Gy (E). Brackets with asterisks compared to 15 Gy: **** = p<0.0001; Money signs compared to 0 Gy: $$$$ = p<0.0001, ns = nonsignificant; WR = WR-1065; N (# of chips) ≥ 3, n (# of MBs) ≥ 120. Drugs are used to treat SGm on day 4 for 30 min prior to and 30 min after radiation and glutathione is analyzed on day 8 for Tempol (F), Edaravone (G), N-acetylcysteine (H), 50 µM Rapamycin (I), 100 ng/mL Palifermin (J), and 50 µM Ex-Rad (K). Brackets with astericks compared to 15 Gy: ns = nonsignificant, **** = p<0.0001; Money sign compared to 0 Gy: $$$$ = p<0.0001; NAC = N-acetylcysteine; N (# of chips) ≥ 3, n (# of MBs) ≥ 150.

Next, the level of senescence was measured using the other reported radioprotective drugs. Tempol (**Fig. 3F**) and Edaravone (**Fig. 3G**) reduced senescence at both 1 mM and 4 mM by 75% and 91% for Tempol, 94% and 113% for Edaravone, at 1 and 4 mM, respectively, versus untreated, irradiated controls. Edaravone also reduced senescence by 96% at 100 µM versus untreated, irradiated controls (**Fig. S4A**). For NAC, 83% and 57% reduction in senescence was observed with 1 mM and 4 mM (**Fig. 3H**) and 82% at 10 mM (**Fig. S4B**) versus untreated, irradiated controls. For drugs with non-antioxidant mechanisms, rapamycin showed complete protection (**Fig. 3I),** whereas Palifermin provided 64% protection (**Fig. 3J)** and Ex-Rad conferred only 45% radioprotection (**Fig. 3K**).

A summary of results from the glutathione and senescence assays shows similar trends for radioprotection (**Table S.5)**. The few differences may result from different mechanisms of action and/or assay targets. Logically, the glutathione assay may be more sensitive to antioxidant function (NAC, Tempol, Edaravone), while the senescence assay may be more appropriate for drugs such as rapamycin, which has anti-senescence properties^58^. These results highlight the trade-offs in developing screening assays and point to the benefit of screening with two assays. Although the assays developed are indirect measures of radiation-induced DNA damage, they nonetheless were validated to detect radiation induce cell-damage and drug radioprotection. Moreover, these assays can be used for *in situ* high-content drug screening with multiple replicates (40-50) per test and enhanced throughput compared to immunohistochemical staining for γH2AX.

### Drug library screening identified several promising radioprotective compounds

The glutathione and senescence assays were used to screen a library of FDA-approved drugs (Selleck Chemicals) at 100 µM. Drugs were first screened using the glutathione assay according to the timeline in **Fig. 2A**. Any compound resulting in statistically similar glutathione levels compared to the 0 Gy control was considered a hit (**Fig. 4**, **orange circles**). Hits with the glutathione assay were then tested with the senescence assay and considered a “double hit” if senescence levels were statistically similar to levels at 0 Gy (**Fig. 4, blue circles**). A list of the 438 drugs screened and relevant statistics are shown in **Table S.3**. Overall, 438 drugs from the library were tested with a hit rate of 5.7%, for a total of 25 double hits **(Fig. 4B**) listed in **Table 1**. While this hit rate is higher than many other drug screening reports (0.1 – 0.3%)^59, 60^, this may be due to the high statistical rigor afforded by the tissue chip format. Additionally, phenotypic screens generally have higher hit rates than target-based screens (> 1%)^61, 62^.

**Figure 4:**
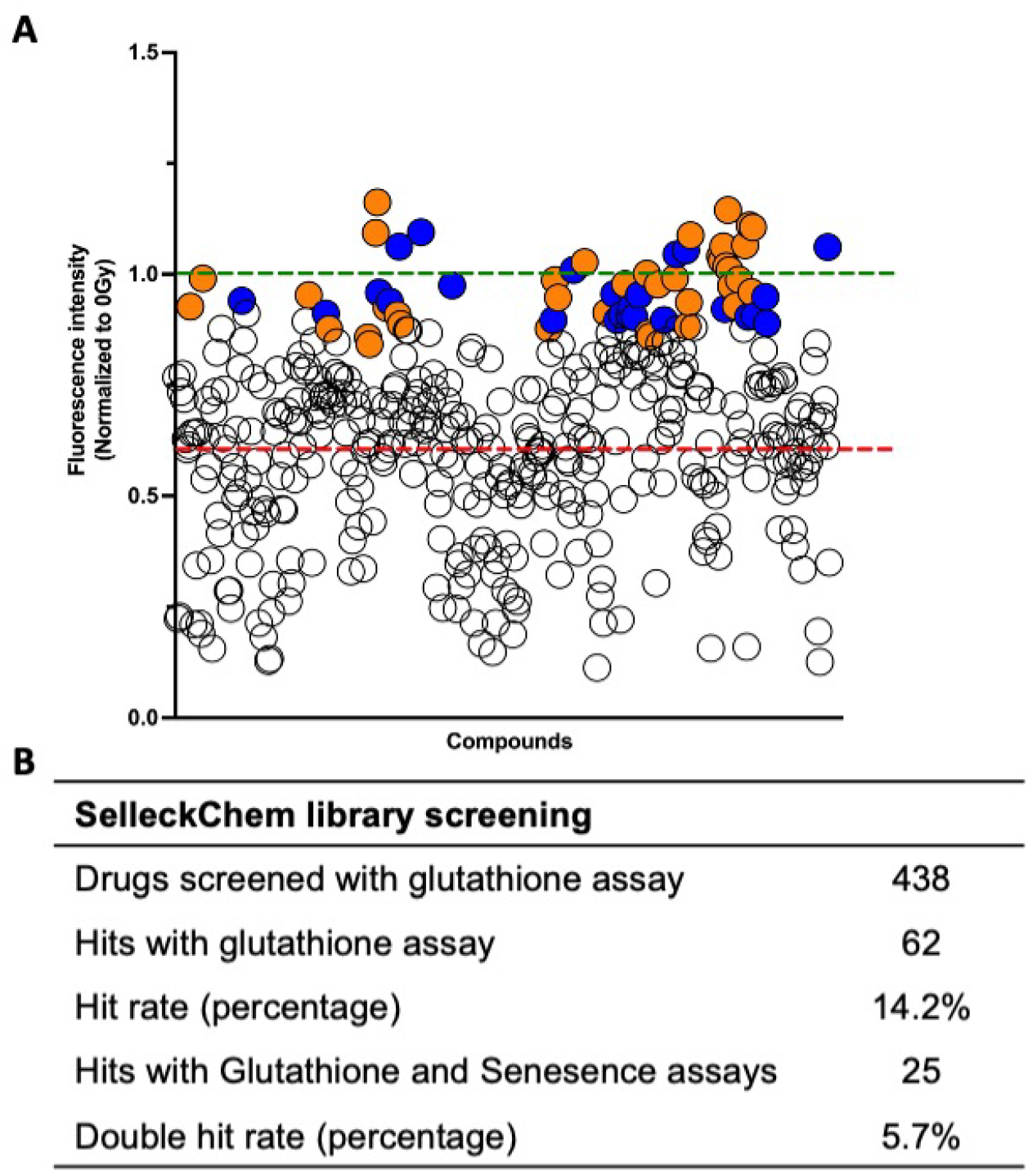
The drug screen identified 25 compounds that protect against post-radiation changes in glutathione and senescence. A) Each circle represents the normalized fluorescence intensity for the glutathione assay for each compound. White circles are compounds that were not hits. Orange circles are compounds that were hits with the glutathione assay only. Blue circles are compounds that were hits with both assays. The green dotted line represents 0 Gy, used for normalization of the data, the red line represents 15 Gy untreated controls using the glutathione assay (A). Table representing the total and percent-positive compounds using the initial glutathione assay and combined glutathione and senescence assays (B).

**Table 1:** Drug activity based on data from the BioAssay database. Drugs highlighted in dark gray were excluded from further analysis due to bioactivity in >10 % of assays. Drugs in light gray were excluded based on poor bioavailability and batch variability (eplerenone)

| Drug | Total Assays | Active (%) |
| --- | --- | --- |
| Glipizide (Glucotrol) | 3141 | 0.5 |
| Etidronate (Didronel) | 1402 | 1.1 |
| Meropenem | 7031 | 1.3 |
| Oxcarbazepine | 2774 | 1.3 |
| Doripenem | 2434 | 1.4 |
| Phenylbutazone | 5565 | 1.5 |
| Enoxacin (Penetrex) | 2673 | 1.9 |
| Cilostazol | 3093 | 2.5 |
| Triamcinolone (Aristocort) | 2345 | 2.7 |
| Didanosine (Videx) | 2840 | 5.7 |
| Eplerenone | 610 | 5.9 |
| Prazosin HCl | 1264 | 6.3 |
| Indomethacin | 10138 | 6.6 |
| Diethylstilbestrol (Stilbestrol) | 5048 | 7.2 |
| Melatonin | 4488 | 8.2 |
| Emtricitabine (Emtriva) | 516 | 9.3 |
| Albendazole (Albenza) | 3086 | 9.3 |
| Rifampin | 8071 | 11.0 |
| Nifedipine (Adalat) | 6190 | 11.5 |
| Rifabutin (Mycobutin) | 1123 | 12.0 |
| Calcitriol (Rocaltrol) | 1786 | 15.2 |
| Lomustine (CeeNU) | 5832 | 16.1 |
| Adefovir Dipivoxil (Preveon) | 929 | 22.7 |
| Curcumin | 3404 | 29.7 |
| Gemcitabine (Gemzar) | 2190 | 46.7 |

### Identification of potential radioprotective drug mechanisms

Of the 25 potential radioprotective compounds, 20 have known interactions with proteins involved in calcium signaling identified within the BioAssay database in PubChem (**Fig. 5A**). These compounds may impact secretory signaling in the salivary gland, which may be radioprotective. While degranulation may not be key to radioprotection, secretory stimulation may play a role in proliferation and survival of the secretory cells^63^. Similarly, using the Drug Set Enrichment Analysis (DSEA) tool to identify pathways, the Reactome analysis related to secretion appear to be upregulated by many of the identified drugs, supporting this potential mechanism (**Fig. 5B,C, and Fig. S5**). Interestingly, only 9 of the 25 compounds have known antioxidant properties, and 12 are anti-inflammatory. This is critical data indicating that alternate mechanisms of radioprotection may be achievable and represented in the identified hits. A reduction in pathway activity related to cell adhesion, cell-cell and cell-matrix interactions also represents a potential area of exploration. Multiple studies have identified changes in integrin expression, cell adhesion and matrix interactions upon radiation exposure which may be linked to cell response^1, 64–67^. Manipulation of these mechanisms in the salivary gland may convey radioprotection.

**Figure 5:**
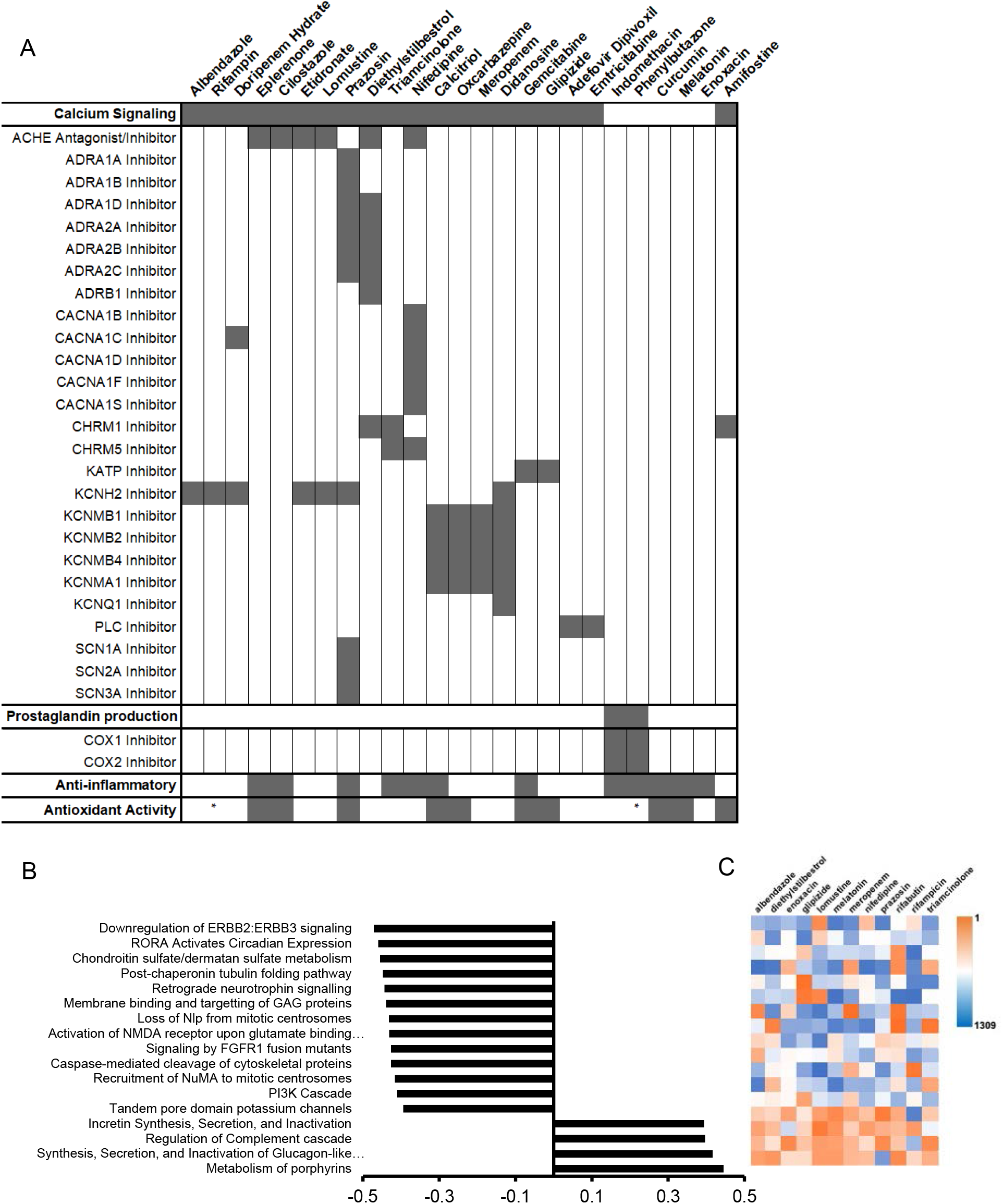
Identification of potential mechanisms of action on salivary gland and similarities among identified radioprotective drugs. Table shows drug activity related to secretion (calcium signaling), prostaglandin production, and anti-inflammatory or antioxidant properties (A)^68–105^. Reactome pathways significantly associated with gene expression patterns affected by treatment with 12 of the identified potential radioprotective compounds (p<0.05). E-scores (−0.5 to 0.5) associated with each pathway show predicted activity related to treatment with the identified drugs (B). Heat map showing ranks for drug-pathway interactions from most-upregulating (orange) to most-downregulating (blue) drug for each pathway (C).

### Hit down-selection using drug promiscuity data and EC_50_ values

A systematic approach was used to down selection the 25 double hits for *in vivo* testing. Since drugs within the library are FDA-approved, considerable information on their pharmacology in mice and humans is readily available through resources such as PubChem. Within PubChem, the BioAssay database was created by the National Institute of Health (NIH) as an open repository containing results of small molecule screening data^38^. We used the BioAssay data to analyze drug promiscuity which refers to the ability of a drug to bind multiple molecular targets with distinct pharmacological outcomes, often causing unwanted side effects^106^. The drugs exhibiting bioactivity in a large number of assays were deprioritized. Data for each double hit was obtained from the database and promiscuity was calculated as the percent of assays reported as “active” (**Table 1**). Drugs with high activity (>10%) were excluded from further testing. Additionally, Etidronate, Melatonin, and Albendazole were excluded due to poor bioavailability^107–109^, and Eplerenone was excluded due to solubility concerns^110^.

The remaining 13 drugs were tested for glutathione levels post-irradiation in dose-limiting experiments (1 −100 µM) to identify effective concentrations. Many drugs were only effective at 100 µM and were excluded due to a lack of potency. For the remaining seven drugs (Phenylbutazone, Meropenem, Diethylstilbestrol, Prazosin, Enoxacin, Glipizide, and Doripenem) dose responses and heat maps summarizing the dose-response results for the glutathione and senescence assays are presented in **Figs. 6 and 7**, respectively. Dose-response curves for the glutathione assay show that Phenylbutazone (**Fig. 6B**) and Meropenem (**Fig. 6C**) exhibited radioprotection equivalent to 0 Gy over a concentration range 0.1 – 100 µM and Diethylstilbestrol (**Fig. 6D**) was radioprotective at 10-100 µM. Prazosin (**Fig. 6E**), Enoxacin (**Fig. 6F**), Glipizide (**Fig. 6G**), and Doripenem (**Fig. 6H**) showed protection only at higher concentrations.

**Figure 6:**
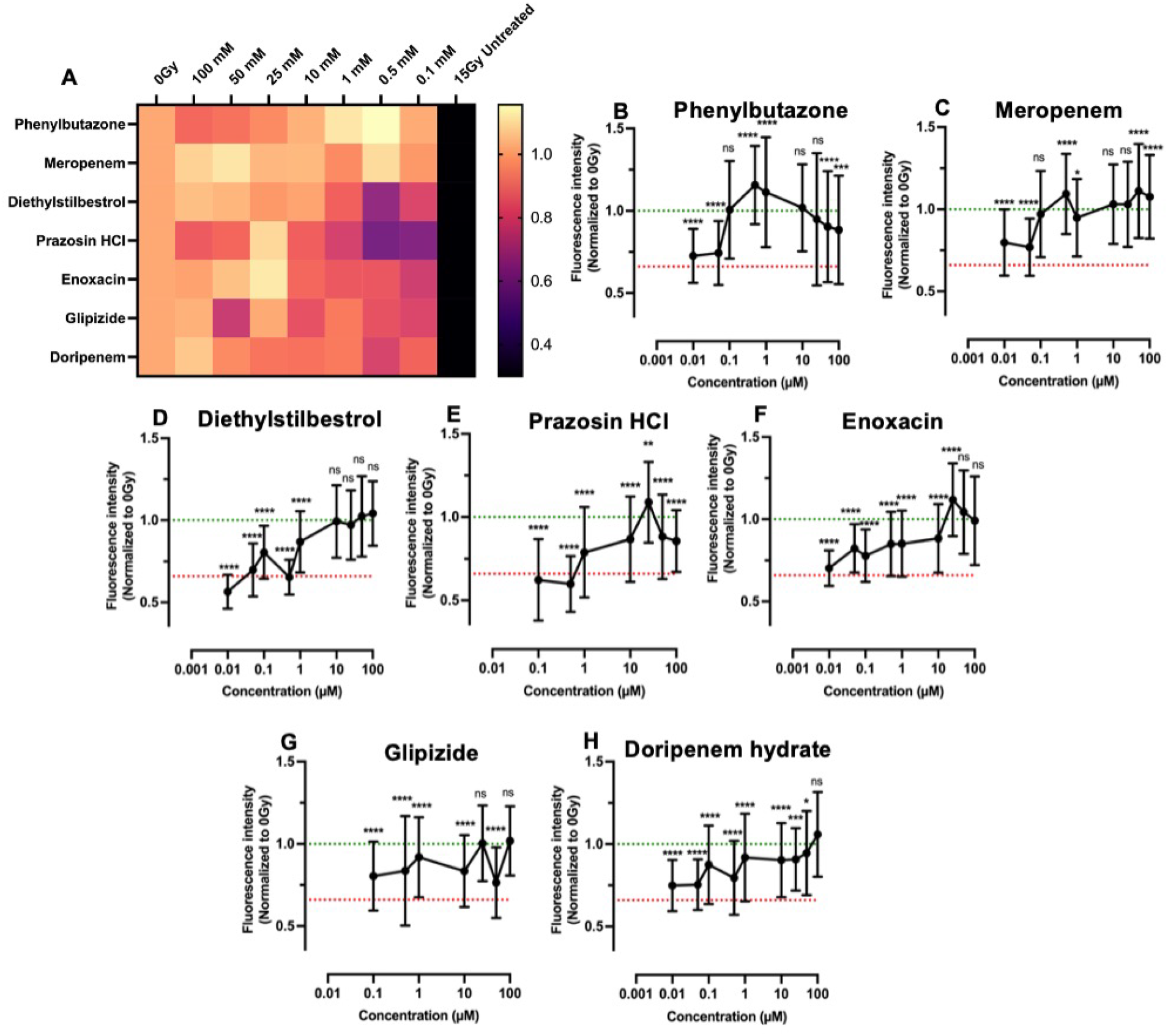
Dose-response data for lead compounds from the drug screen analyzed via glutathione levels. Heat map of combined dose-response results with 0 Gy (1.0) at left and 15 Gy (0.3) only on right. (A). Results for Phenylbutazone (B), Meropenem (C), Diethylstilbestrol (D), Prazosin HCl (E), Enoxacin (F), Glipizide (G), and Doripenem hydrate (H). Green and red lines represent 0 Gy and 15 Gy averages, respectively. Statistics were calculated using ANOVA with Dunnett’s post-hoc test. ns = nonsignificant; **p < 0.01, ***p < 0.001, ****p < 0.0001

**Figure 7:**
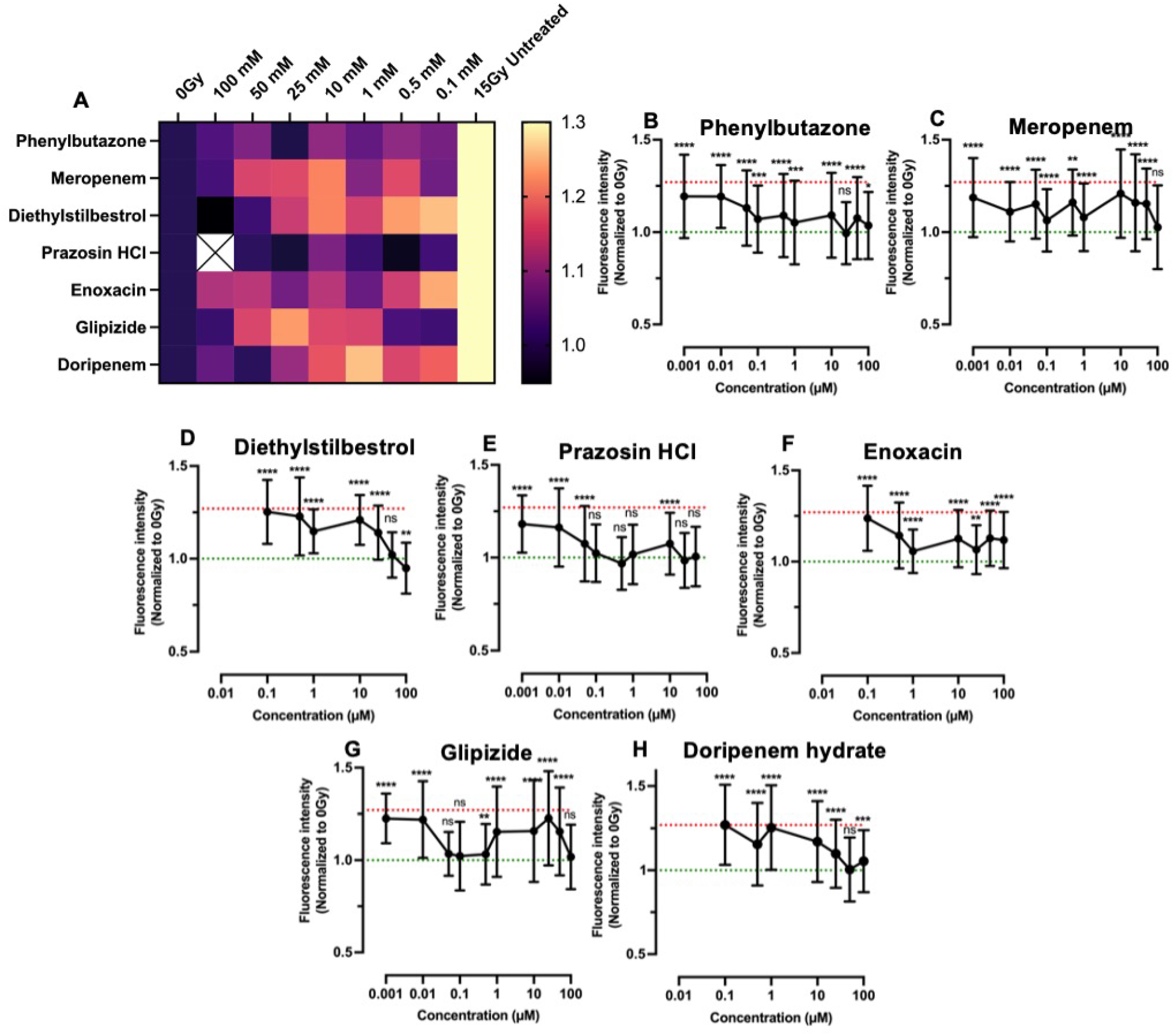
Dose-response data for lead compounds from the drug screen analyzed for senescence levels. Heat map of combined dose-response results with 0 Gy (1.0) at left and 15 Gy (1.3) only on right. (A). Results for Phenylbutazone (A), Meropenem (B), Diethylstilbestrol (C), Prazosin HCl (D), Enoxacin (E), Glipizide (F), and Doripenem hydrate (G). Green and red lines represent 0 Gy and 15 Gy averages, respectively. Statistics were calculated using ANOVA with Dunnett’s post-hoc test. ns = nonsignificant; **p < 0.01, ***p < 0.001, ****p < 0.0001

The radioprotection trends based on the senescence assay differed somewhat, with Phenylbutazone (**Fig. 7B**) and Meropenem (**Fig. 7C**) showing only partial protection and Diethylstilbestrol (**Fig. 7D**) exhibiting radioprotection equivalent to 0 Gy between 50-100 µM concentrations whereas Prazosin (**Fig. 7E**) showed complete protection between 0.1-100 µM. Glipizide (**Fig. 7G**) and Doripenem (**Fig. 7H**) also showed variable protection.

EC_50_ values extrapolated from dose response curves are shown in **Table 2**. Phenylbutazone showed the most promising results, with low EC_50_ values for both the glutathione (0.08 µM) and senescence (0.05 µM) assays. Phenylbutazone is a non-steroidal anti-inflammatory drug (NSAID) that inhibits cyclooxygenases (COX-1 and COX-2), enzymes that produce prostaglandins^111^. Prostaglandins, specifically PGE_2_ signaling, have been shown to increase in irradiated salivary glands, and mitigation of salivary gland damage was achieved through treatment with the anti-inflammatory drug Indomethacin^1, 112^. Indomethacin also showed radioprotection in our drug screen but was ineffective at concentrations lower than 100 µM. Indomethacin only blocks COX-1, underscoring the greater efficacy of Phenylbutazone. Phenylbutazone was originally developed for chronic pain for conditions such as arthritis but has since been restricted to treating ankylosing spondylitis due to induction of rare but severe blood disorders, including anemia and leukopenia^111^. However, doses ranged from 300-1000 mg, generating a plasma concentration of 30-50 µg/mL. In contrast, the EC_50_ of 0.08 µM for protecting against radiation-induced glutathione changes established in this study equates to a 26 ng/mL concentration. Thus, the risk of severe adverse effects may be greatly diminished for doses necessary for radioprotection. Additionally, Phenylbutazone has excellent bioavailability (up to 90%) ^111, 113^ and long half-life (50-105 hrs) ^111^, which may enable dose de-escalation, further decreasing risks.

**Table 2:** Calculated EC_50_ values for the top radioprotective drugs. Values were estimated using a nonlinear fit in Prism. “Undetermined” indicates that the software could not accurately fit a curve to the data.

| Compound | Estimated EC<sub>50</sub> ($\mu\text{M}$) | |
| --- | --- | --- |
|  | Glutathione | Senescence |
| Phenylbutazone | 0.08 | 0.05 |
| Meropenem | 0.1 | undetermined |
| Diethylstilbestrol | 0.7 | 37.7 |
| Prazosin HCl | 1 | 0.04 |
| Enoxacin | 2.4 | 0.5 |
| Glipizide | 4.0 | undetermined |
| Doripenem hydrate | 8.6 | 16.4 |

Enoxacin, an antibacterial agent used for treating urinary tract infections^114^, was previously identified as radioprotective^115^. Using a high-throughput screening method with the viability of lymphocytes as the primary readout, two classes of antibiotics (tetracyclines and fluoroquinolones) were identified as robust radioprotectors, including Enoxacin ^115^. This observation corroborates our drug screening results, in which several antibiotics were identified as double hits, including Enoxacin, Meropenem, Doripenem hydrate, Rifampin, and Rifabutin. The Enoxacin EC_50_ of 2.4 µM for the glutathione assay is similar to the 13 µM EC_50_ reported for viability of lymphocyte cells^115^. Notably, five other fluoroquinolones reported as radioprotectors (Levofloxacin, Gatifloxacin, Ofloxacin, Moxifloxacin, and Norfloxacin)^115^ were not hits in our drug screen (**Table S3**). These disparities may be related to differences in cell type (salivary gland versus lymphocyte) or readouts (glutathione/senescence versus viability).

Based upon drug down selection data and measured EC_50_ values, Phenylbutazone, Enoxacin, and Doripenem hydrate were selected for *in vivo* validation in mice with γH2AX foci per nucleus IHC staining as an outcome measure consistent with prior work showing correlation with the development of xerostomia^17, 41, 42, 46^. Vehicle-treated SGm exposed to 15 Gy radiation exhibited a 3.3-fold increase in the number of γH2AX foci per nucleus, indicating a significant increase in double-stranded DNA breaks due to radiation exposure (**Figs. 8A-D, I**). Treatment with WR-1065 resulted in a 0.5-fold reduction in γH2AX foci per nucleus compared to 15 Gy controls (**Figs. 8D, E, I**). No significant differences existed between 0 Gy controls and WR-1065 treated SGm exposed to 15 Gy (**Figs. 8C, E, I**). These results are similar to prior studies utilizing WR-1065 *in vitro* and *in vivo* via retrograde ductal injection^17, 42^. Treatment with the test compounds, Phenylbutazone, and Enoxacin resulted in 0.4- and 0.5-fold reduction in γH2AX foci per nucleus compared to 15 Gy controls, respectively **(Figs. 8E-G, I**). Results observed after Phenylbutazone treatment were not significantly different from 0 Gy controls (**Figs. 8C, F, I)**. Enoxacin treatment resulted in a 1.6-fold increase in γH2AX foci per nucleus **(Figs. 8C, G, I).** Treatment with Doripenem hydrate did not reduce γH2AX foci per nucleus relative to 15 Gy controls and showed a 2.7-fold increase compared to 0 Gy controls **(Figs. 8C, D, H, I).**

**Figure 8:**
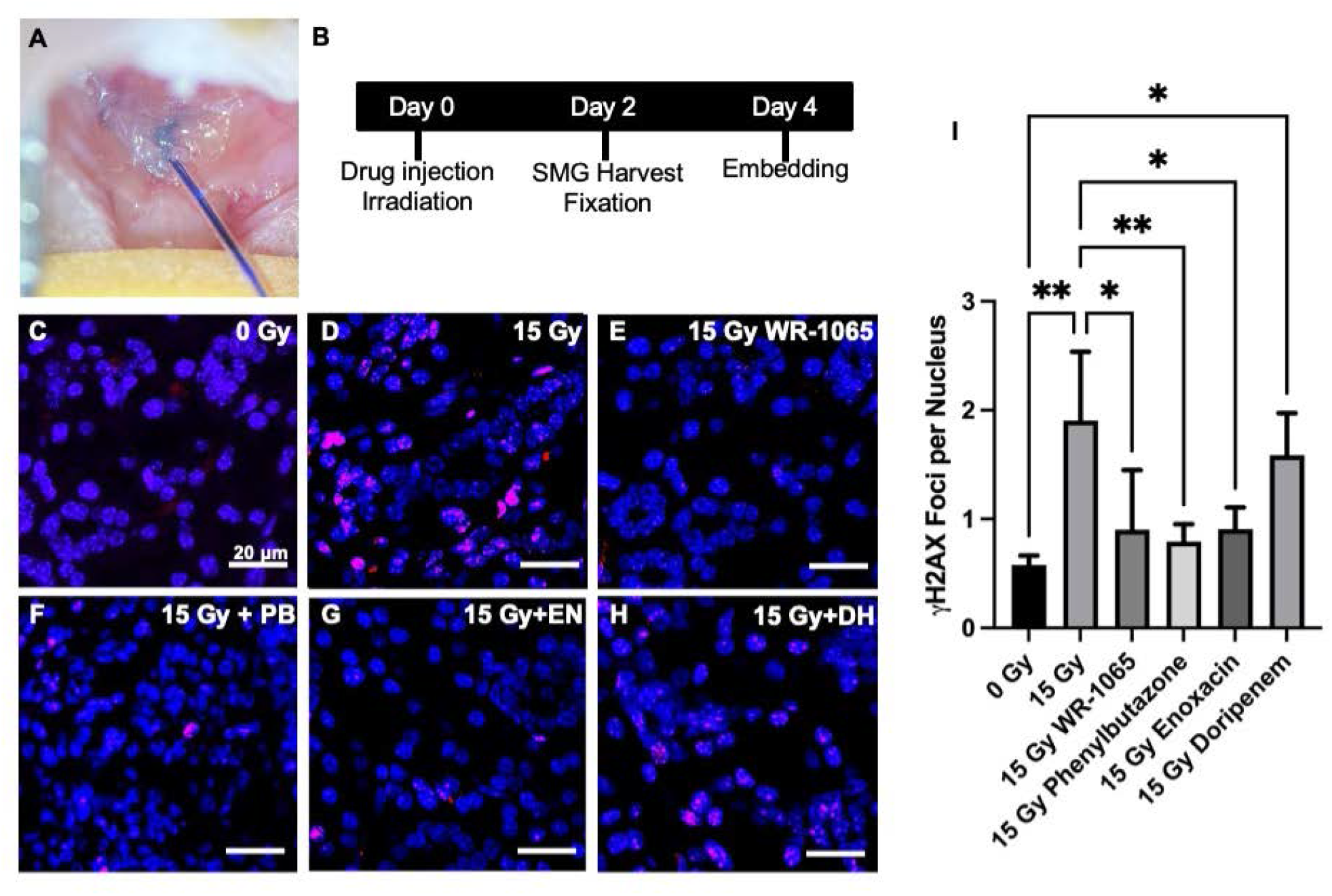
*In vivo* testing of double hits for radioprotection of mouse salivary gland. Representative image of retroductal drug administration (A). Schematic of drug treatment, irradiation, and harvest protocol (B). Representative images of γH2AX staining in mouse salivary gland tissue 48 hours post treatment with vehicle (C) and 15Gy radiation exposure (D), as well treatment with control drug WR1065 (E) and lead drug hits, Phenylbutazone (PB) (F), Enoxacin (EN) (G), and Doripenem hydrate (DH) (H) followed by 15 Gy radiation exposure. Scale bars: 20 μm. Quantification of γH2AX foci per nucleus from all treatment groups (I). (N=4-6)* p<0.05, ** p<0.01 vs. 0 Gy, # p<0.05, ## p<0.01 vs. 15 Gy.

## 4. Conclusions

To overcome long-standing challenges associated with off-target radiation damage resulting in life-long dry mouth and poor quality of life, we sought to leverage our salivary gland tissue chip technology^17^ to identify novel radioprotective drugs. First, we investigated several assays that detect cell damage following radiation exposure (**Table S1**). We identified reduced glutathione and β-galactosidase (cell senescence) as biomarkers of radiation damage and developed assay protocols for high-content screening. We validated the assays with known radioprotective drugs, including WR1065, the active metabolite of Amifostine, the only FDA-approved preventative therapy for radiation-induced xerostomia^14, 15, 51^. We next tested other drugs reported to be radioprotective including Tempol^20, 21, 49^, Edaravone^23, 56, 57^, N-acetylcysteine^22^, Palifermin^27, 28^, Ex-Rad^25, 26^, and Rapamycin^24^. Our results found that Tempol and Edaravone showed complete radioprotection similar to WR1065 by both assay whereas the other drugs showed partial radioprotection. Partial protection likely reflects in differences in tissue types or the assay protocols used in the other studies but our results confirm that all the drugs tested exhibit radioprotection at some concentration, by one or both assays (**Table S5**). A library of FDA-approved drugs was screened, enabling the de novo identification of 25 drugs with radioprotective activity (**Table 1**). Of the 25 hits, 20 drugs have known interactions with proteins involved in calcium signaling (**Fig. 5A**), which is critical for saliva secretion^116^. Only 9 of the 25 compounds have known antioxidant properties, and 12 are anti-inflammatory. This suggests alternate mechanisms of radioprotection. A DSEA pathways analysis suggests these drugs upregulate secretion and activate pathways related to cell adhesion, cell-cell and cell-matrix interactions suggesting possible new mechanistic targets for designing novel radioprotective drugs. Using the PubChem database and experimentally-determined EC_50_ values (Figs. 6, 7 **and Table 2)**, the list of 25 double hits was narrowed down to 3 candidates (Phenylbutazone, Enoxacin, and Doripenem) for testing using an *in vivo* radiation damage model. In vivo, results demonstrate that Phenylbutazone and Enoxacin have equivalent radioprotection to Amifostine (**Fig. 8)**.

In conclusion, this body of work validates the SGm tissue chip as a high-content drug discovery tool. We validated glutathione and β-galactosidase (cell senescence) as biomarkers of ionizing radiation damage. Novel drugs were discovered that confer radioprotection by mechanisms other than antioxidant activity. Ongoing studies seek to investigate chemical analogs and mechanisms of action to identify lead drug candidates that can be developed for clinical translation.

## Acknowledgements

The authors gratefully acknowledge the support of the National Institute of Dental and Craniofacial Research (NIDCR), and the National Center for Advancing Translational Sciences (NCATS) of the National Institutes of Health, under award numbers UG3DE027695 and UH3 DE002212 to DSWB, CEO, and LAD, T32 ES007026 to JAM, and F31 DE029658 to LRP. The content is solely the responsibility of the authors and does not necessarily represent the official views of the National Institutes of Health. The authors would like to thank Matthew Ingalls for helpful discussions.

## 5. Supplemental Materials

**Figure S1:**
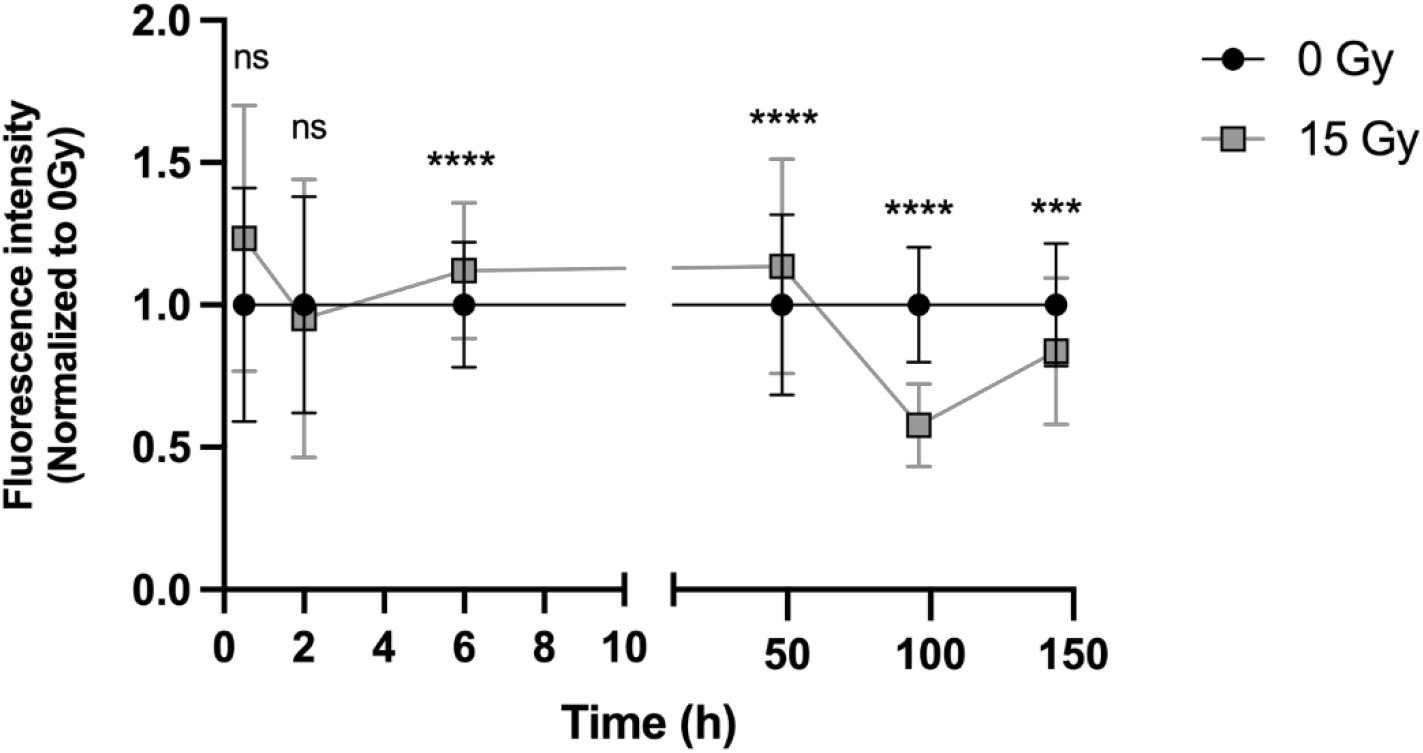
The reduced glutathione assay shows the best signal separation at 4 days post-radiation. Fluorescence intensity is normalized to 0 Gy. Statistics are comparing 0 Gy to 15 Gy at each timepoint. ns = nonsignificant, ****p < 0.0001, ***p < 0.001, N (# of chips) ≥ 3, n (# of MBs) ≥ 120. I’m surprised that there is significance except 96 h. Not great support of specificity of glutathione.

**Figure S2:**
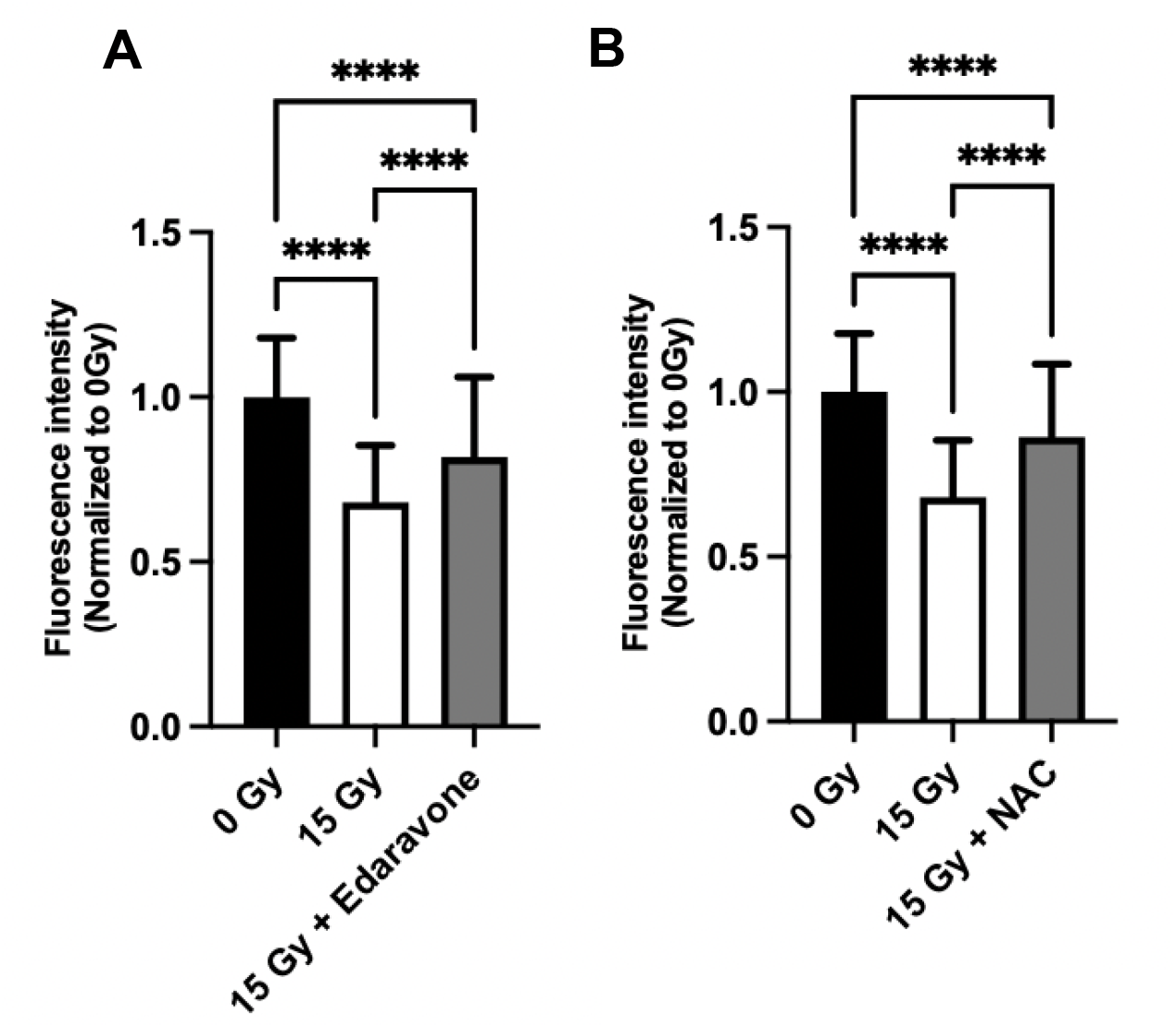
Edaravone and N-acetylcysteine show partial radioprotection using the glutathione assays. Edaravone (A) tested at 0.1 mM and N-acetylcysteine (B) tested at 10 mM. ****p < 0.0001; NAC = N-acetylcysteine; N (# of chips) ≥ 3, n (# of MBs) ≥ 200.

**Figure S3:**
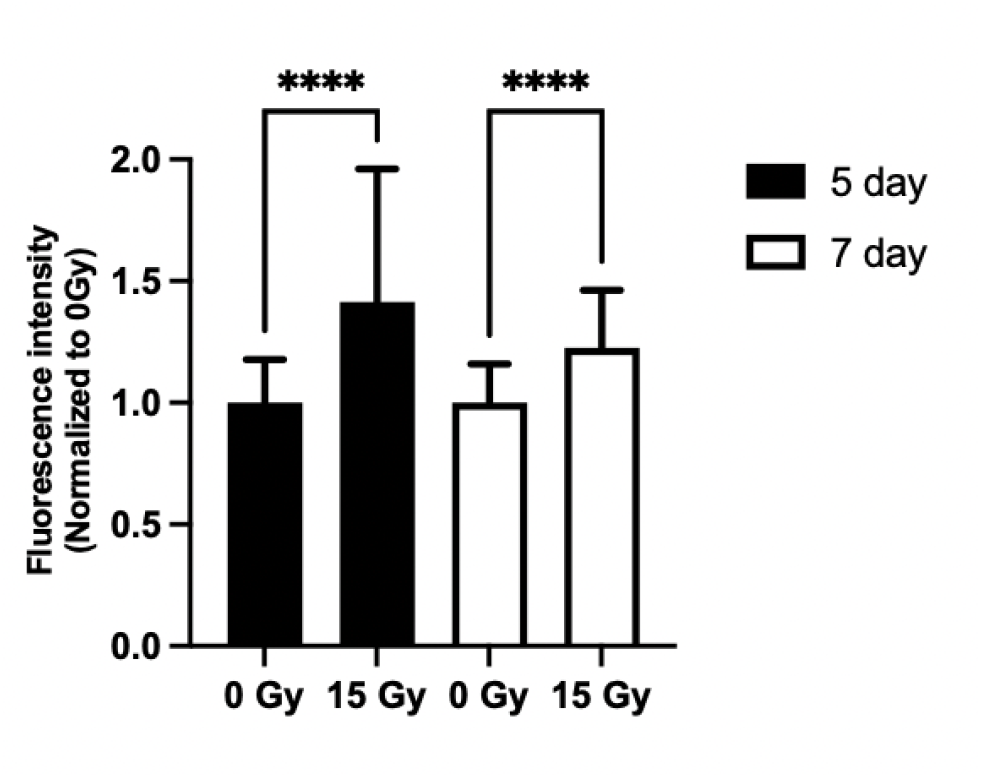
The β-Galactosidase senescence assay shows the greatest signal separation at 5 days post-radiation. Senescence assay tested at 5 and 7 days. N ****p < 0.0001; N (# of chips) ≥ 4, n (# of MBs) ≥ 180.

**Figure S4:**
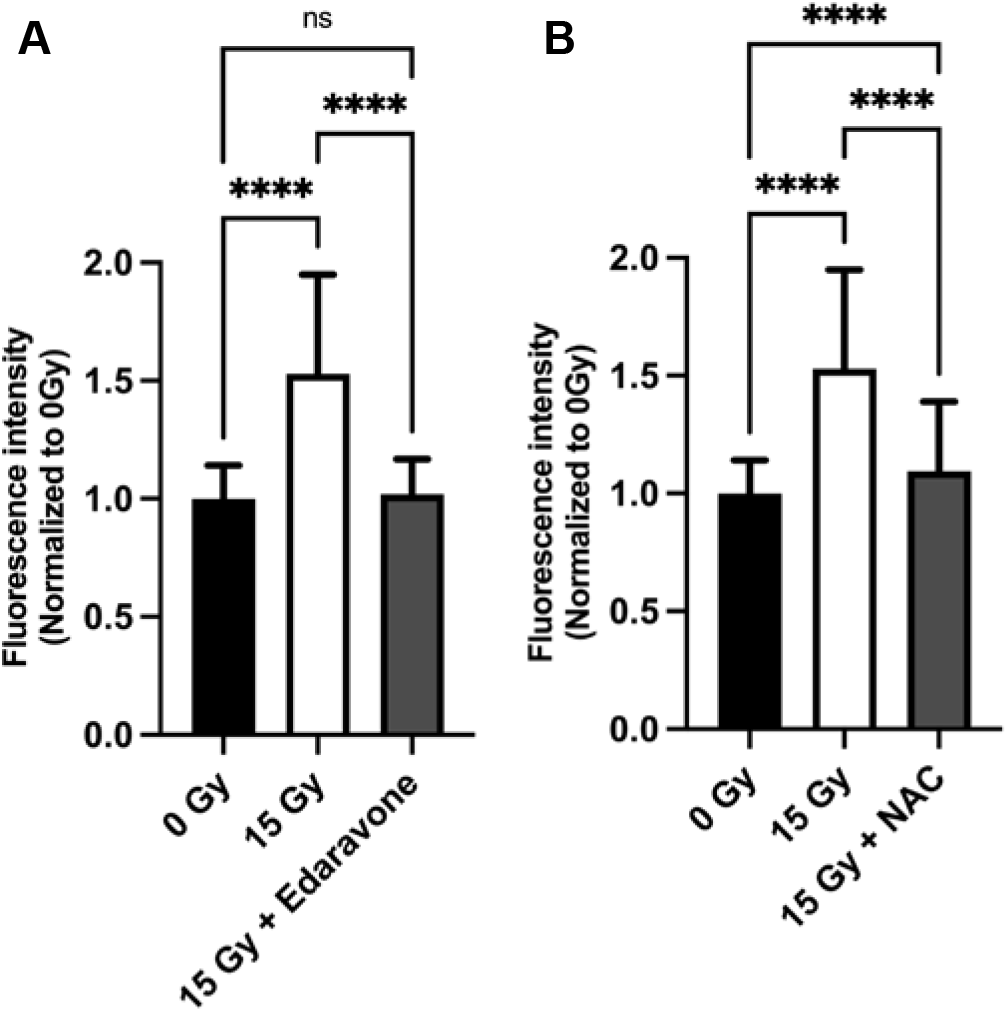
Edaravone and N-acetylcysteine are protective against radiation-induced senescence ay high concentrations. Edaravone (A) tested at 0.1 mM and N-acetylcysteine (B) tested at 10 mM. ****p < 0.0001; NAC = N-acetylcysteine; N (# of chips) = 3, n (# of MBs) ≥ 120.

**Supplemental Figure S5:**
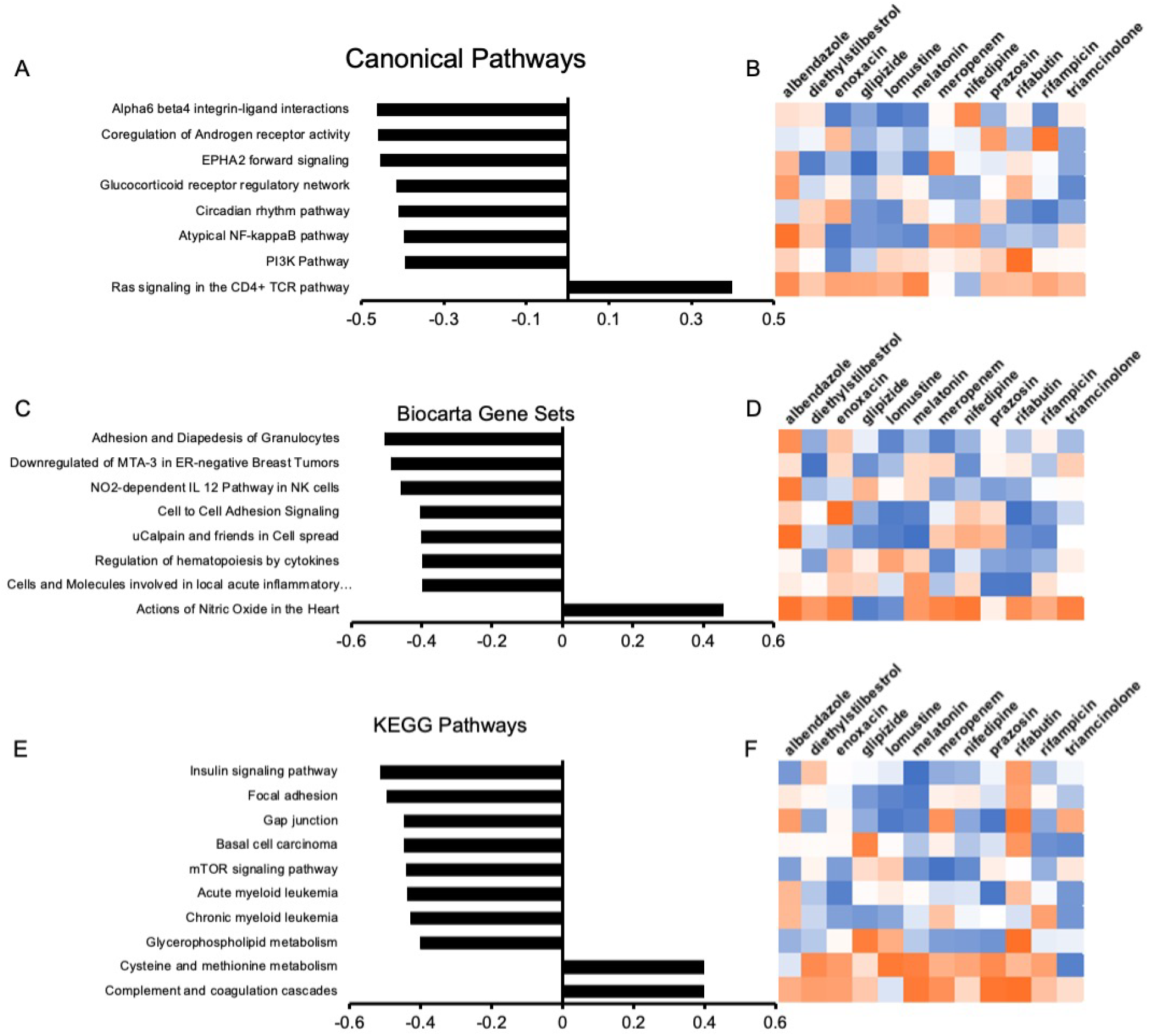
Identification of potential drug mechanisms and similarities among identified radioprotective compounds. E-scores associated with each pathway show predicted activity related to treatment with the identified compounds A) canonical pathways, C) Biocarta gene sets, and E) KEGG pathways. Heat maps showing ranks for drug-pathway interactions from most-upregulating (orange) to most-downregulating (blue) drug for each pathway from B) canonical pathways, D) Biocarta gene sets, and F) KEGG pathways.

**Supplemental Table S.1:**
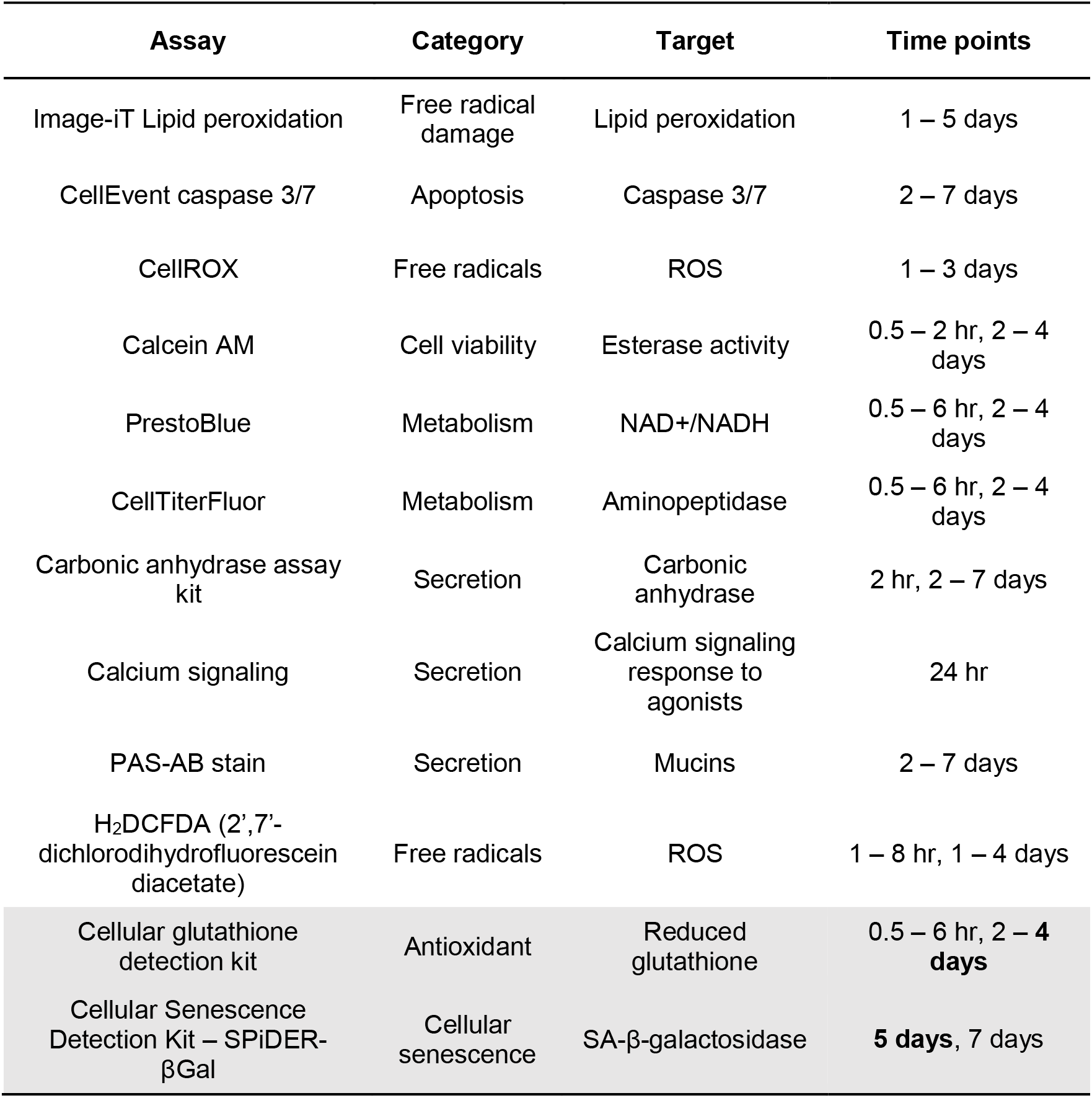
Assays tested for detecting radioprotection in the salivary gland tissue chip. Rows in gray are the assays capable of detecting a reliable change between 0 Gy and 15 Gy at the time points in bold.

| Assay | Category | Target | Time points |
| --- | --- | --- | --- |
| Image-iT Lipid peroxidation | Free radical damage | Lipid peroxidation | 1 – 5 days |
| CellEvent caspase 3/7 | Apoptosis | Caspase 3/7 | 2 – 7 days |
| CellROX | Free radicals | ROS | 1 – 3 days |
| Calcein AM | Cell viability | Esterase activity | 0.5 – 2 hr, 2 – 4 days |
| PrestoBlue | Metabolism | NAD<sup>+</sup>/NADH | 0.5 – 6 hr, 2 – 4 days |
| CellTiterFluor | Metabolism | Aminopeptidase | 0.5 – 6 hr, 2 – 4 days |
| Carbonic anhydrase assay kit | Secretion | Carbonic anhydrase | 2 hr, 2 – 7 days |
| Calcium signaling | Secretion | Calcium signaling response to agonists | 24 hr |
| PAS-AB stain | Secretion | Mucins | 2 – 7 days |
| H<sub>2</sub>DCFDA (2',7'-dichlorodihydrofluorescein diacetate) | Free radicals | ROS | 1 – 8 hr, 1 – 4 days |
| Cellular glutathione detection kit | Antioxidant | Reduced glutathione | 0.5 – 6 hr, 2 – **4 days** |
| Cellular Senescence Detection Kit – SPiDER-βGal | Cellular senescence | SA-β-galactosidase | **5 days**, 7 days |

**Table S.2:** Product information for radioprotective drugs used to validate the glutathione and senescence assays

| **Drug** | **Vendor** | **Cat. #** | **Solvent** | **Storage** |
| --- | --- | --- | --- | --- |
| **4-Hydroxy-TEMPO (Tempol)** | Sigma Aldrich | 176141 | DI water | Stored at 4 °C; fresh solution mixed each time |
| **N-acetylcysteine** | Sigma Aldrich | A9165 | DI water | Stored at 4 °C; fresh solution mixed each time |
| **3-Methyl-1-phenyl-2-pyrazoline-5-one (Edaravone)** | Sigma Aldrich | M70800 | DMSO | Stored at room temp, fresh solution mixed each time |
| **WR1065** | Sigma Aldrich | W2020 | DI water | -20 °C |
| **Rapamycin** | MedChemExpress | HY-10219 | DMSO | -80 °C |
| **Ex-Rad (Recilisib)** | MedChemExpress | HY-101625 | DMSO | -80 °C |
| **Keratinocyte Growth Factor (Palifermin)** | ThermoFisher (Invitrogen) | PHG0094 | PBS | -20 °C |

**Table S.3:**
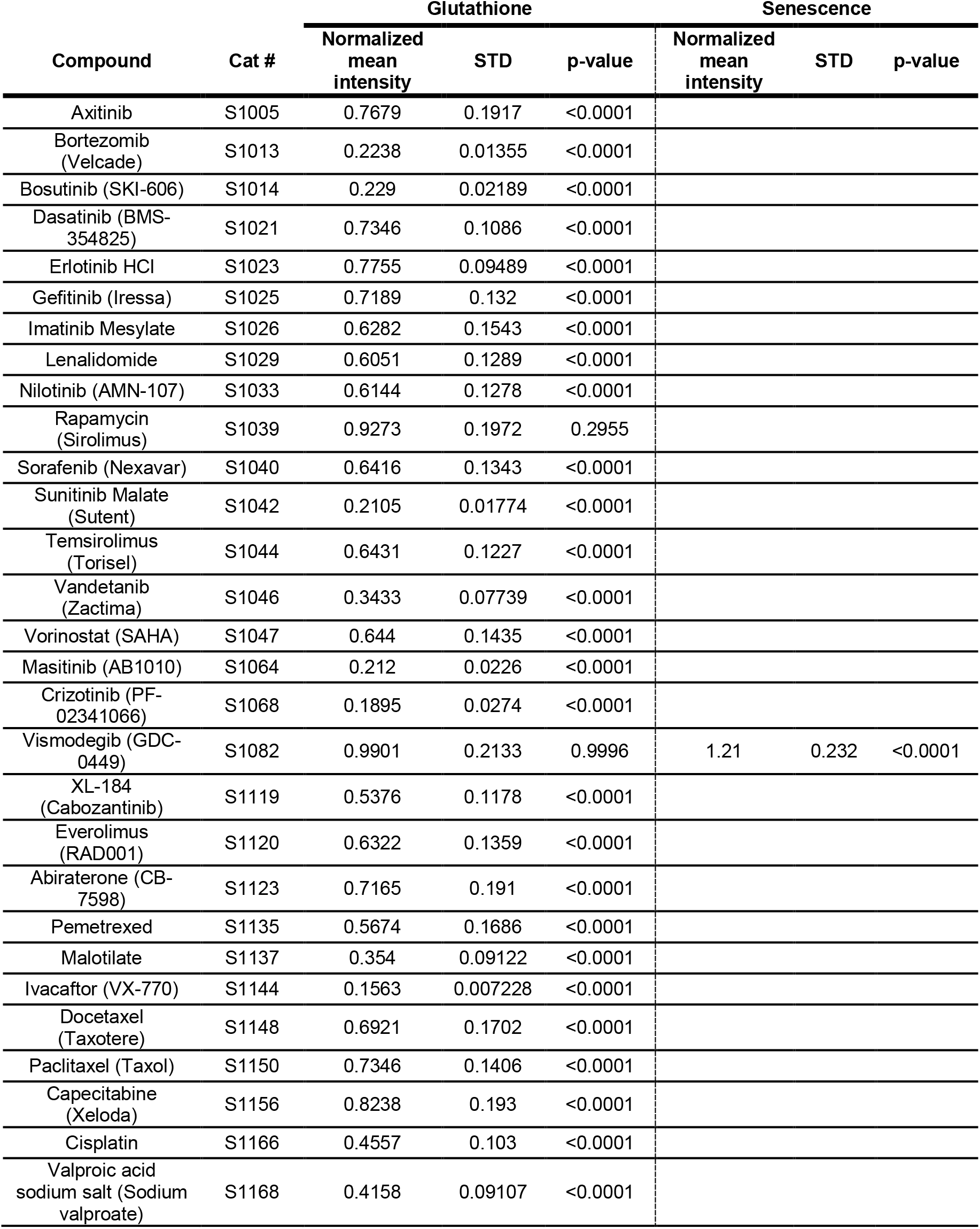
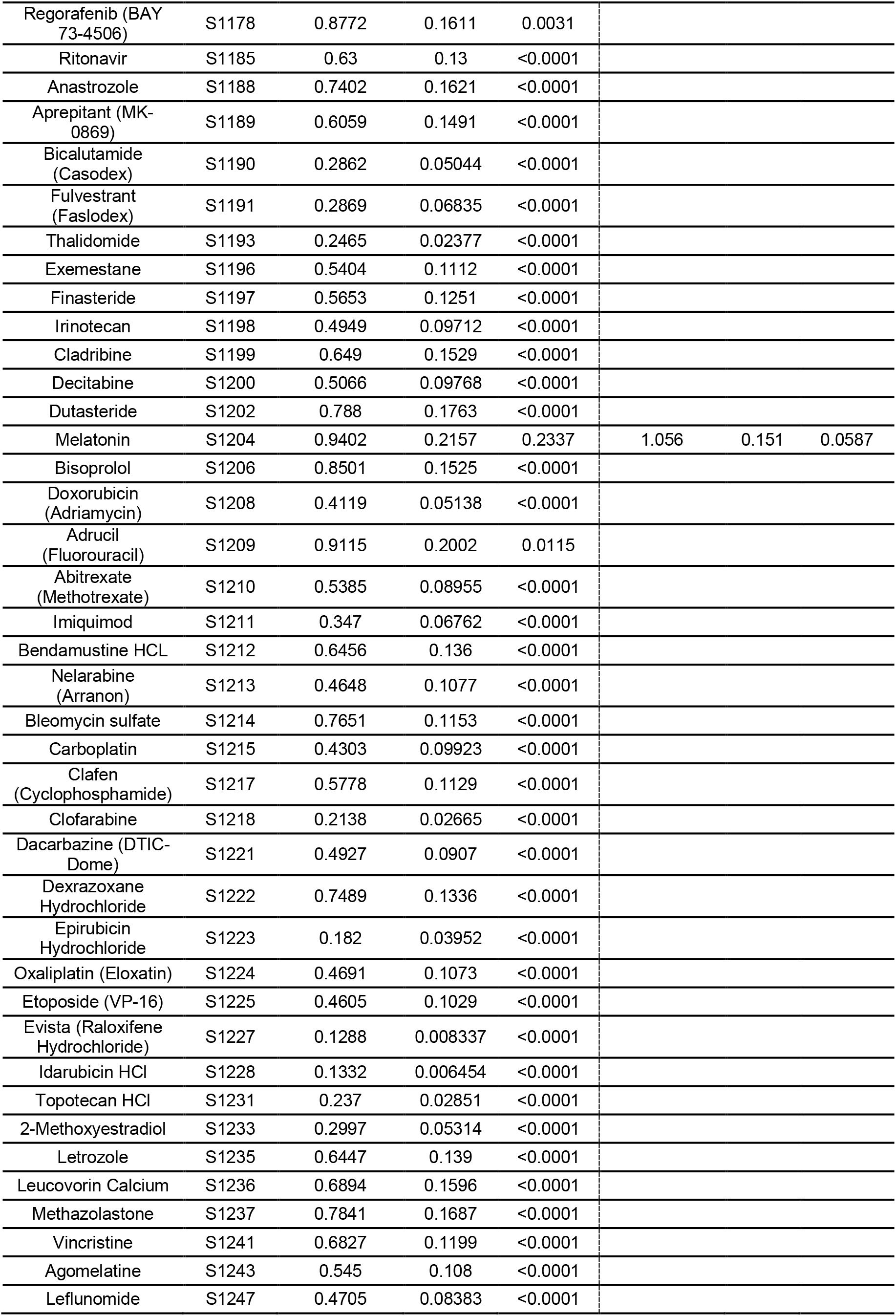
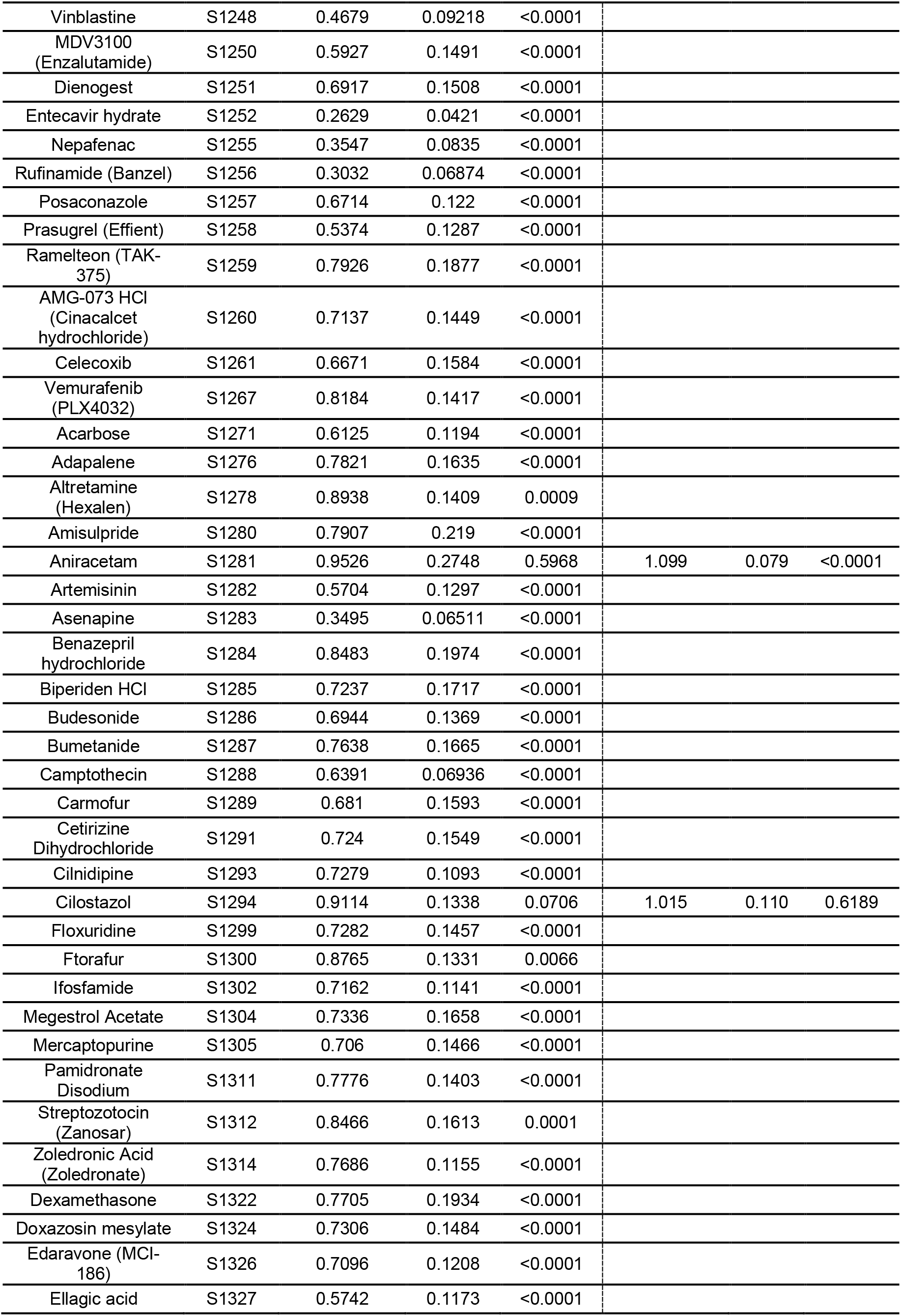
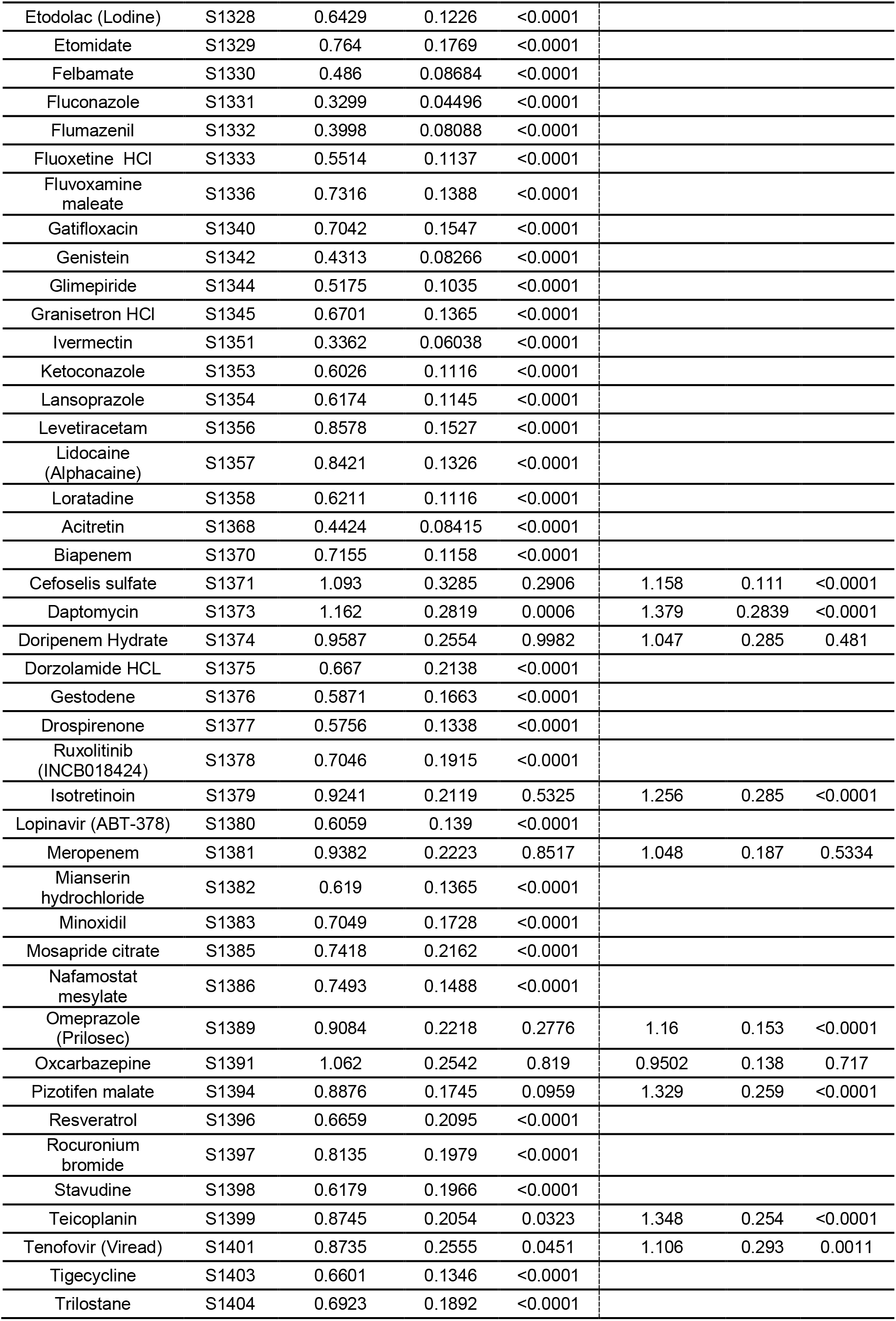
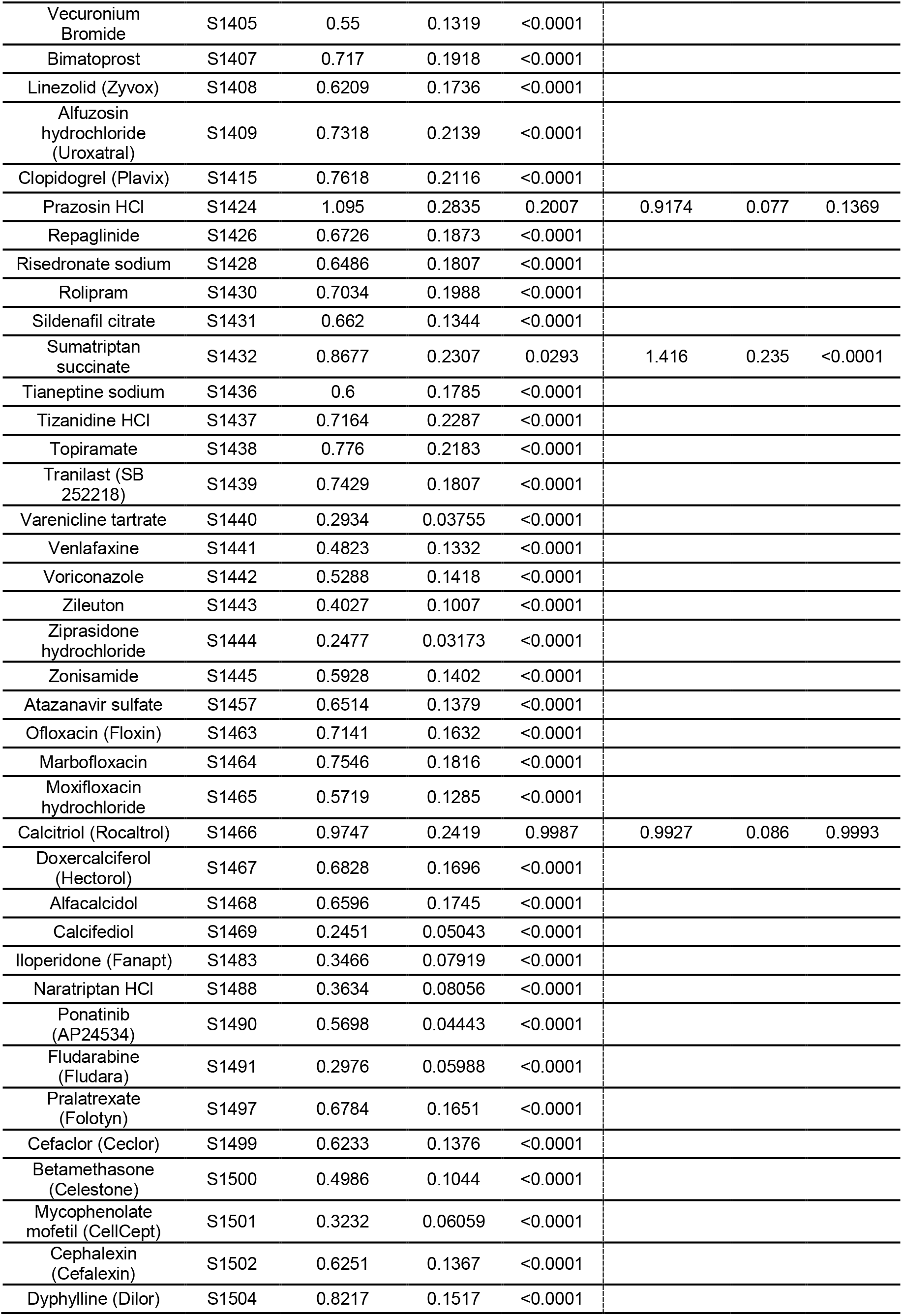
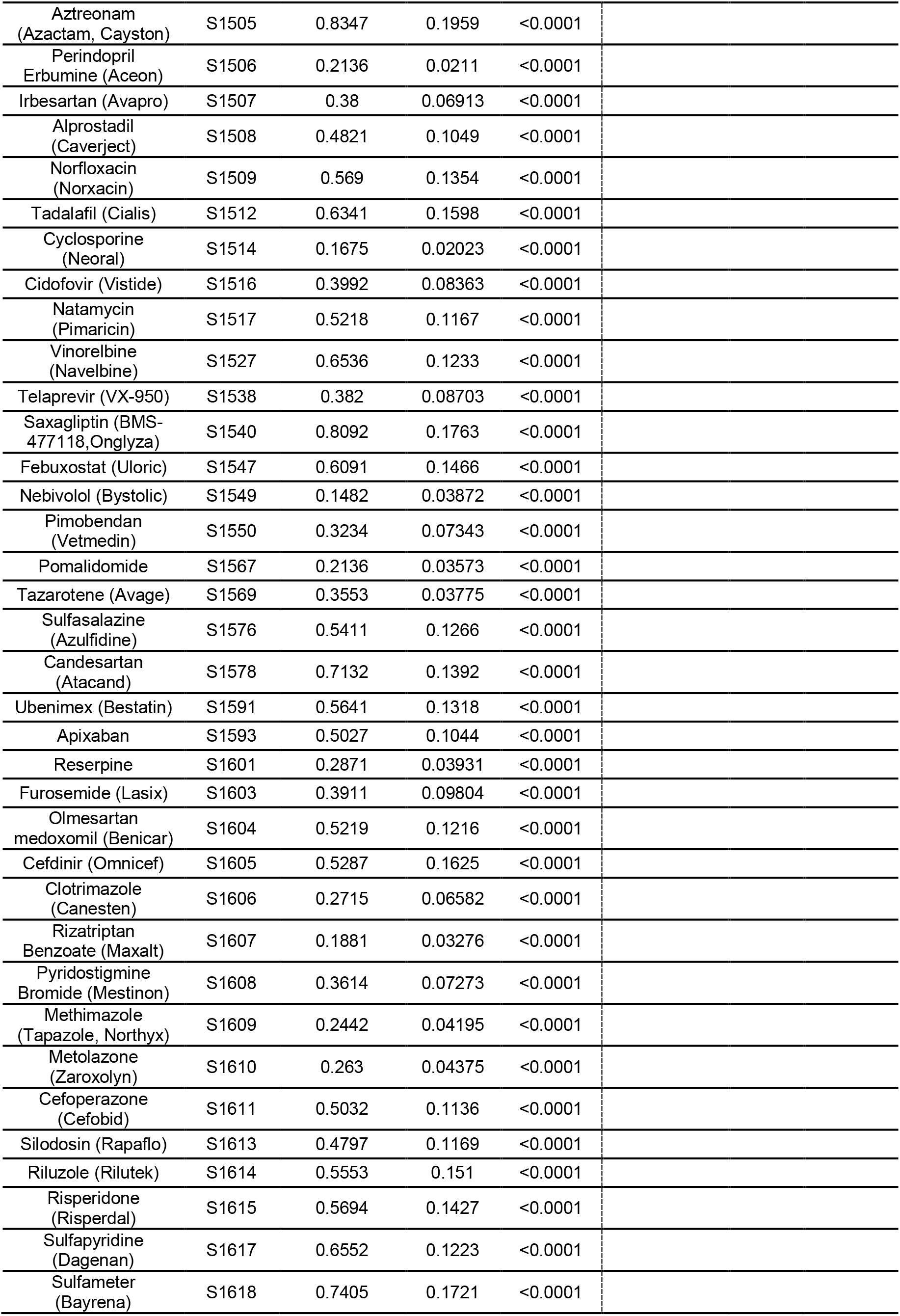
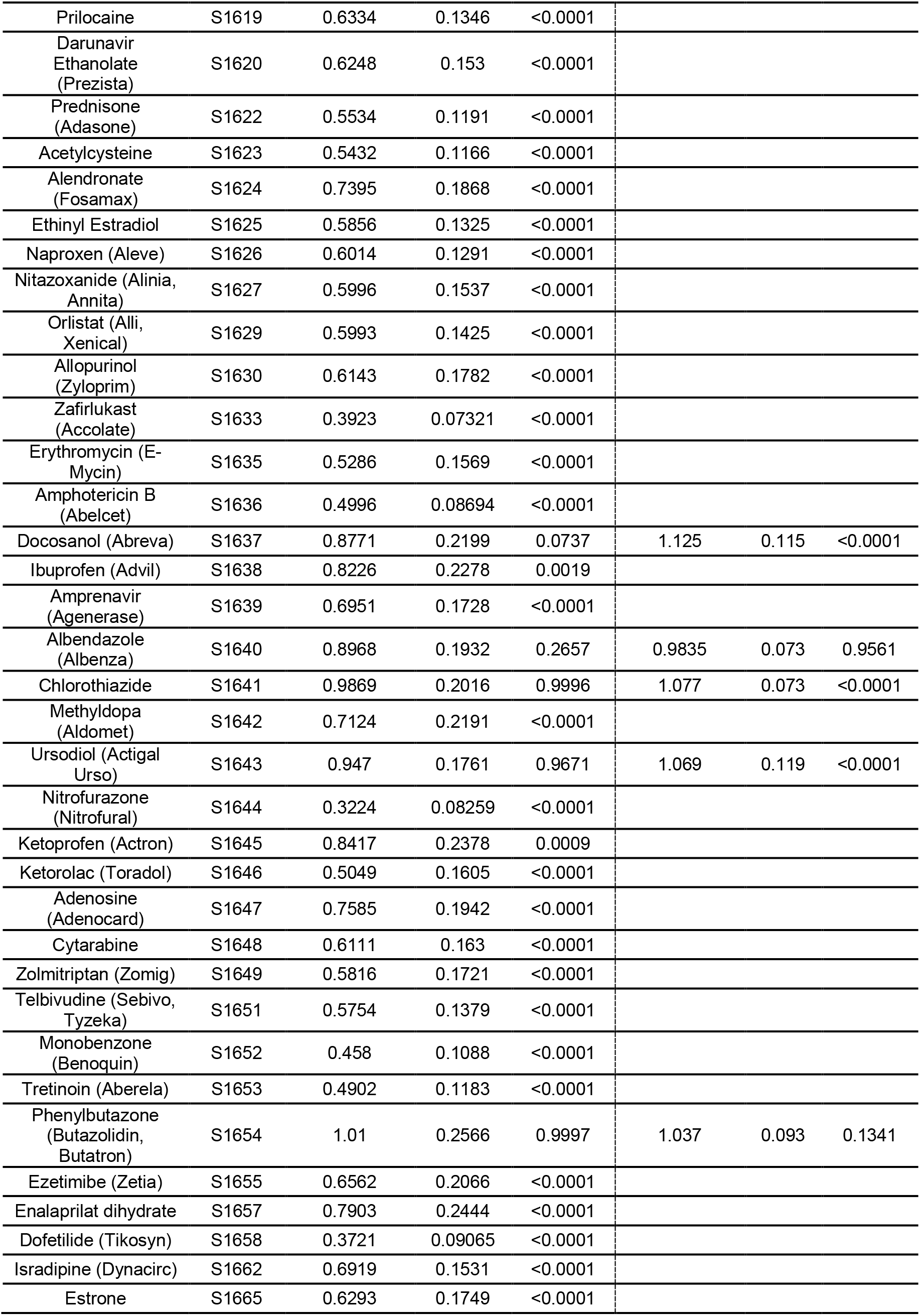
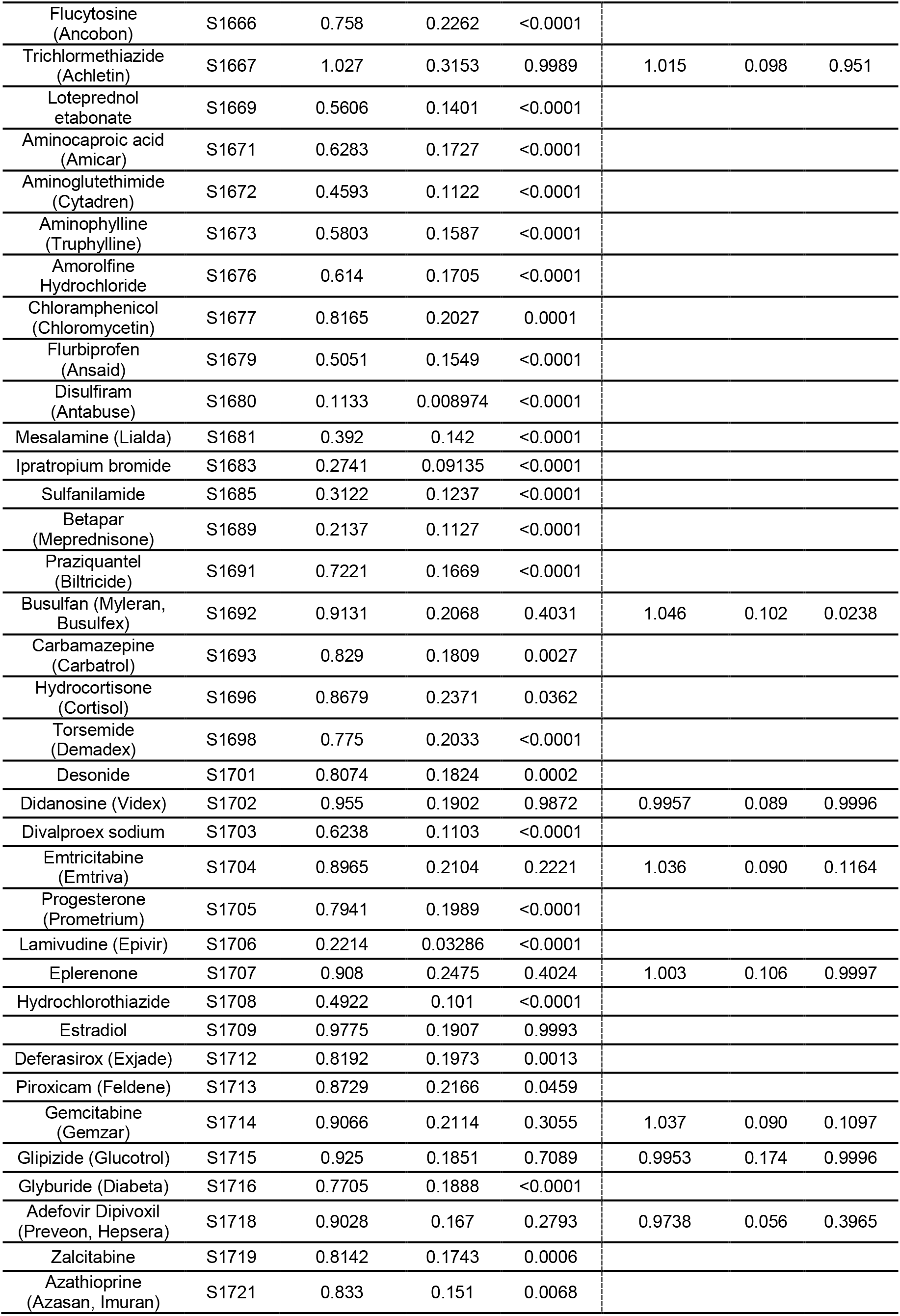
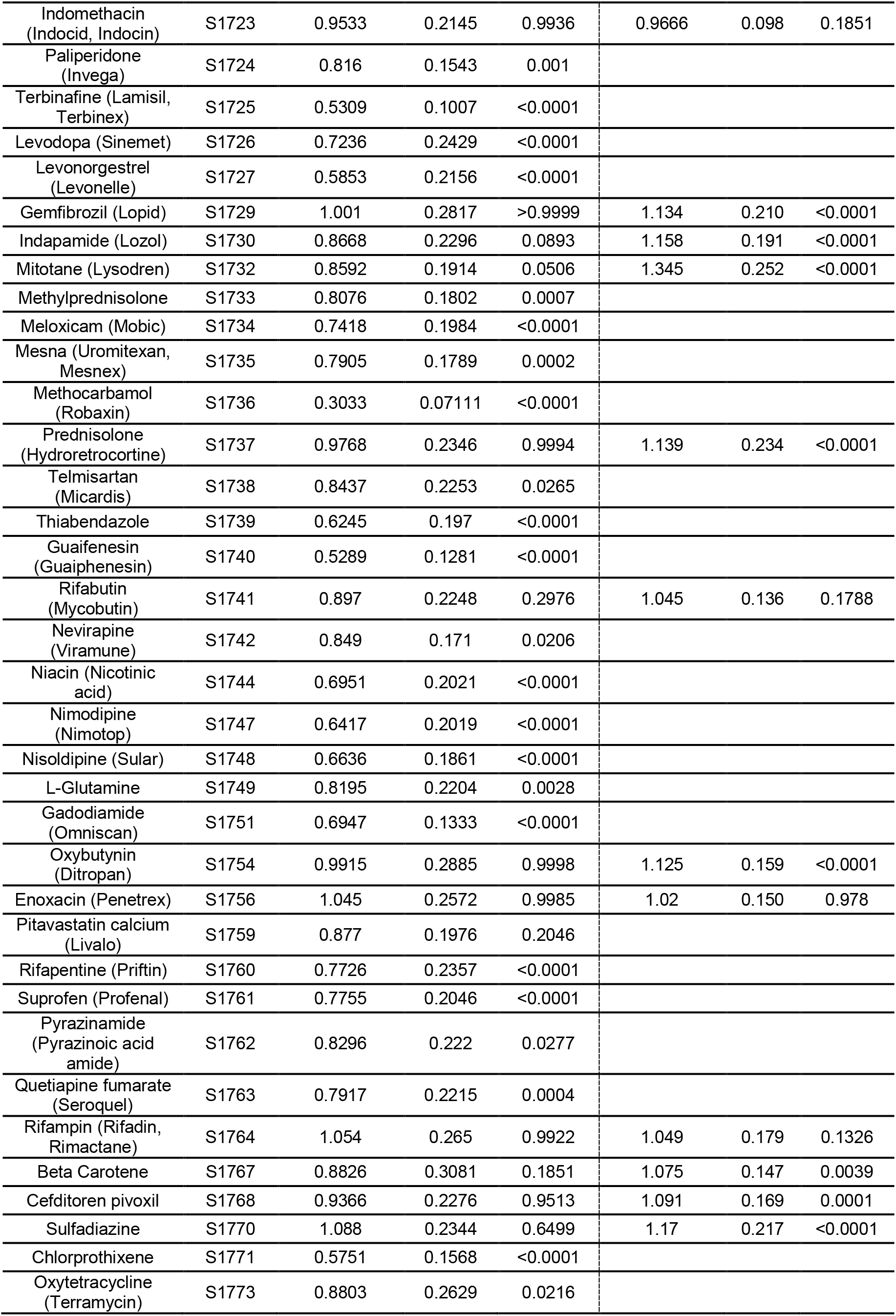
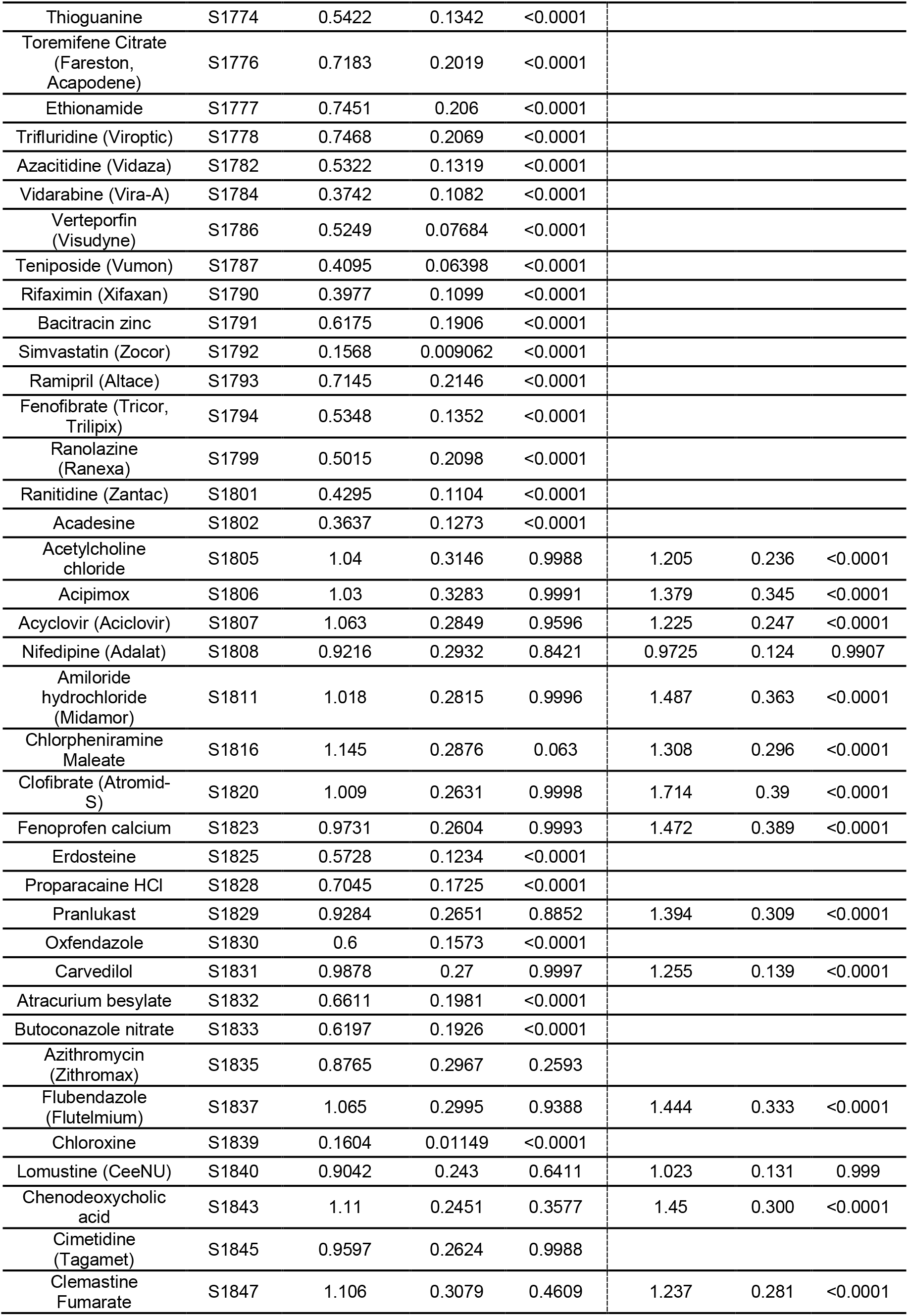
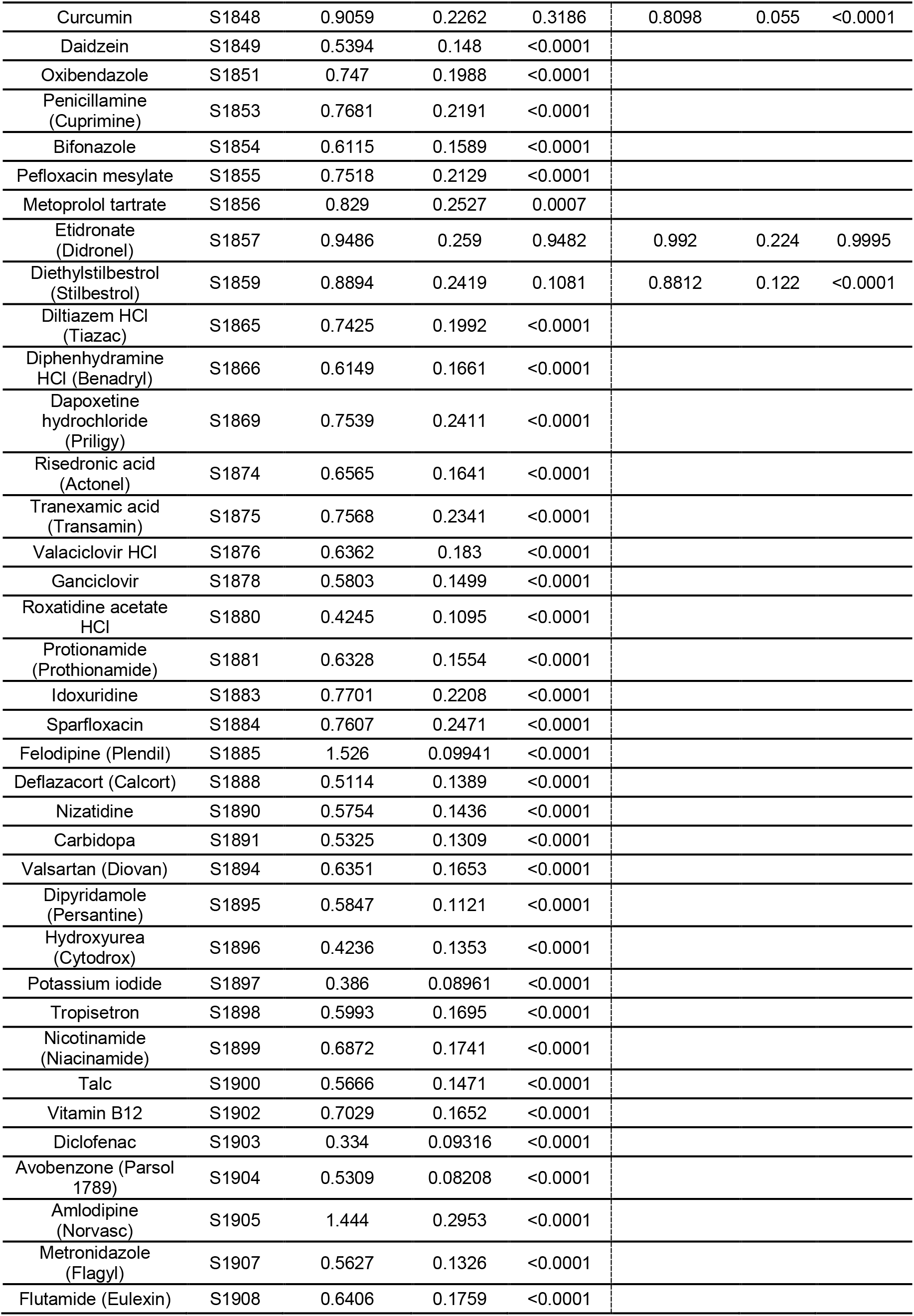
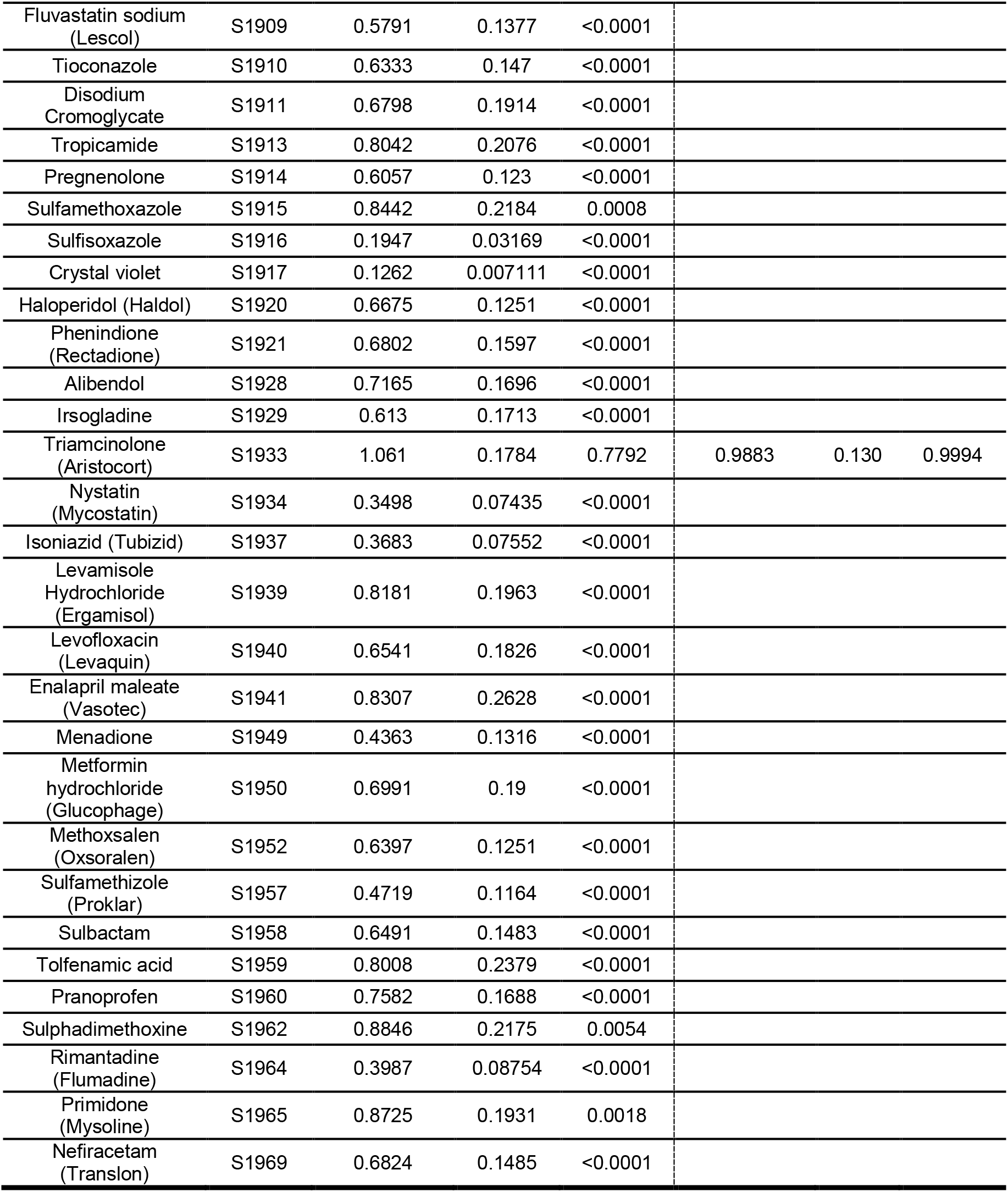
List of compounds tested from the Selleck Chemicals drug library. The normalized mean intensity, standard deviation (STD), and p-values compared to 0 Gy are shown for each assay. Only drugs that were nonsignificant compared to 0 Gy in the glutathione assay were tested with the senescence assay.

**Table S.4:**
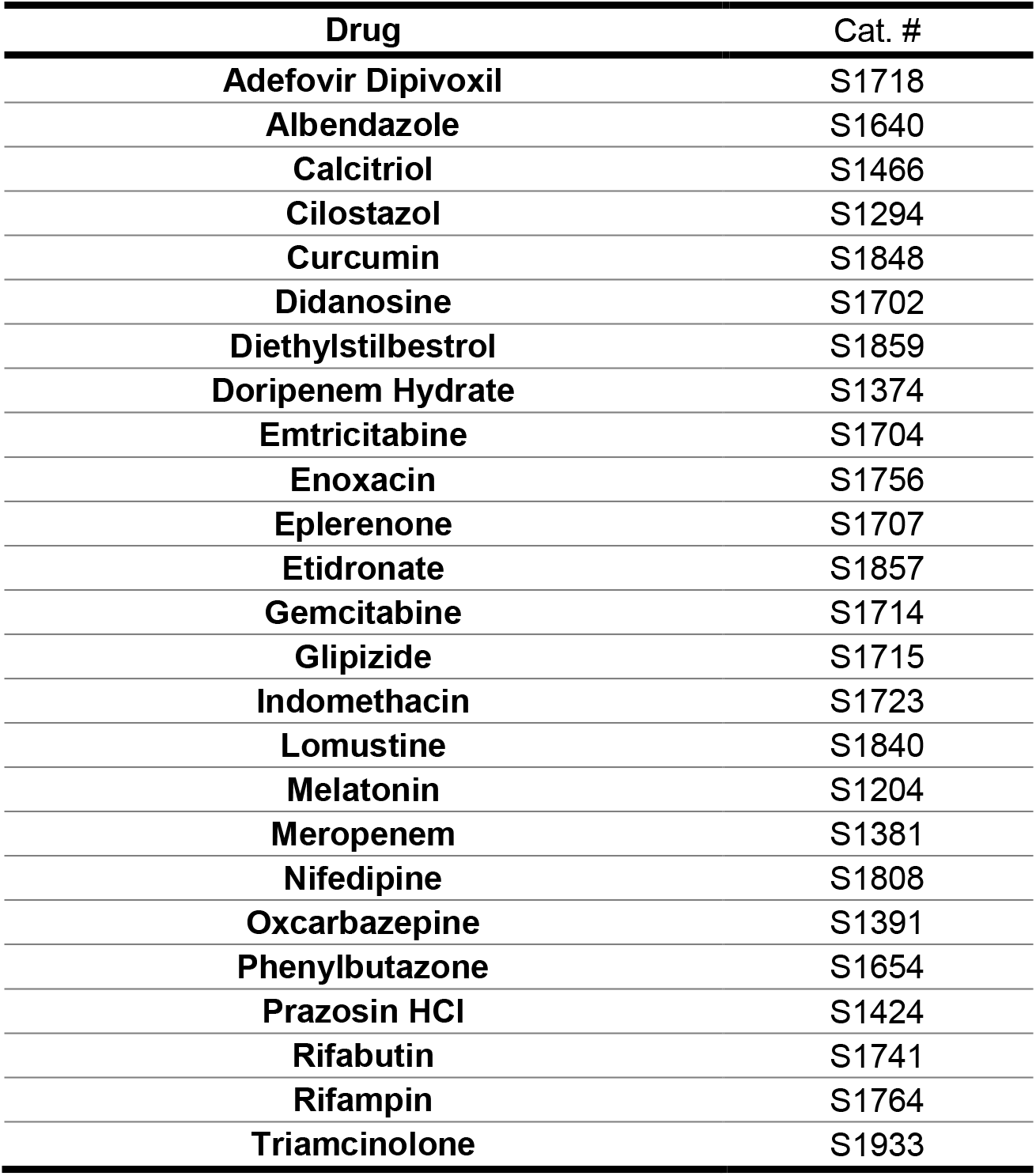
Drugs identified as double hits in the drug screen.

**Table S.5:** Summary of the results of known radioprotective drugs tested with the glutathione and senescence assays.

| Drug | Concentration | Glutathione | Senescence |
| --- | --- | --- | --- |
| WR-1065 | 1 mM | ✓ | ✓ |
|  | 4 mM | ✓ | ✓ |
| Tempol | 1 mM | ✓ | ✓ |
|  | 4 mM | ✓ | ✓ |
| Edaravone | 100 µM | ~ | ✓ |
|  | 1 mM | ✓ | ✓ |
|  | 4 mM | ✗ | ✓ |
| N-acetylcysteine | 1 mM | ✗ | ~ |
|  | 4 mM | ✗ | ~ |
|  | 10 mM | ~ | ~ |
| Rapamycin | 50 µM | ~ | ✓ |
| Palifermin | 100 ng/mL | ~ | ~ |
| Ex-Rad | 50 µM | ✗ | ~ |

✓ = radioprotection; ✗ = no radioprotection; ~ = partial radioprotection

